# Spatio-temporal dynamics of nuclear CREB1: what does it mean?

**DOI:** 10.1101/2022.06.26.497665

**Authors:** Luz E. Farias Altamirano, Elena Vásquez, Carlos L. Freites, Jorge E. Ibañez, Mario E. Guido, Estela M. Muñoz

**Affiliations:** Instituto de Histología y Embriología de Mendoza (IHEM), Universidad Nacional de Cuyo (UNCuyo), Consejo Nacional de Investigaciones Científicas y Técnicas (CONICET), Mendoza, Argentina; CIQUIBIC-CONICET, Departamento de Química Biológica “Ranwel Caputto”, Facultad de Ciencias Químicas, Universidad Nacional de Córdoba, Córdoba, Argentina

**Keywords:** bZIP transcription factor family, cyclic AMP responsive element-binding protein 1 (CREB1), phosphorylation, pineal gland (PG), spatio-temporal dynamics, cellular heterogeneity, superior cervical ganglionectomy (SCGx), RNA polymerase II

## Abstract

In the mammalian pineal gland (PG), cyclic AMP responsive element-binding protein 1 (CREB1) participates in the nocturnal melatonin synthesis that rhythmically modulates physiology and behavior. Phosphorylation of CREB1 present in pinealocyte nuclei is one of the key regulatory steps that drives pineal transcription. The spatio-temporal dynamics of CREB1 itself within PG cell types have not yet been documented. In this study we analyzed total CREB1 via Western blot, and the dynamism of CREB1 nuclear distribution in individual rat pinealocytes using fluorescence immunohistochemistry followed by confocal laser-scanning microscopy and quantitative analysis. Total CREB1 levels remained constant in the PG throughout the light:dark cycle. The distribution pattern of nuclear CREB1 did vary, however, among different PG cells. Pinealocytes emerged as having discrete CREB1 domains within their nucleoplasm that were especially distinct. The number, size, and location of CREB1 foci fluctuated among pinealocytes, within the same PG and among *Zeitgeber* times. A significantly larger dispersion of CREB1-immunoreactive nuclear sites was found at night. This was not accompanied by changes in the overall transcription activity, which was mostly conserved between the light and dark phases, as shown by the expression of a particular phosphorylated form of the RNA polymerase II (RNAPII-pSer^5^CTD). Suppression of the nocturnal norepinephrine pulse by chronic bilateral superior cervical ganglionectomy increased CREB1 dispersion in pinealocyte nuclei, as compared to sham-derived cells. In addition, differences in CREB1 distribution were found between sham-operated and non-operated rats at early night. Together, these data suggest that in mature pinealocytes nuclear CREB1 is subjected to a dynamic spatio-temporal distribution. Further studies are necessary to elucidate the underlying mechanisms, including the role of chromatin and interchromatin elements, and to understand the impact of CREB1 reorganization in the pineal transcriptome.

## Introduction

In vertebrates, the pineal gland (PG) is part of a multicentric circuit that imposes circadian rhythmicity to physiology and behavior (1, 2). Nowadays, it is widely accepted that melatonin, produced by pinealocytes at night, serves as the indole hormone responsible for disseminating this chronobiological cue (3). The circadian production of melatonin is a highly conserved phenomenon among vertebrate species, and its underlying molecular mechanisms are well understood (4, 5). In rat, *de novo* gene expression is required, and the activating transcription factor (TF) CREB1 [cyclic AMP (cAMP) responsive element-binding protein 1] and the inhibiting TF ICER (inducible cAMP early repressor; an isoform of the cAMP responsive element modulator, CREM) are key regulators of the melatonin rhythm (6–8). CREB1 and ICER coordinate with many other transcription actors to finely shape rhythmic melatonin synthesis (2, 4, 9–11).

More generally, CREB is a prototypical stimulus-inducible TF that is mainly located in the nuclear compartment of the cell (7, 8, 12–14). This 43-kDa basic leucine zipper (bZIP) member of the CREB/CREM/ATF-1 (activation transcription factor 1) family couples gene expression to a wide spectrum of extracellular stimuli and intracellular signals, including cAMP, Ca^2+^ and cytokines (14–17).

CREB is widely expressed across many multicellular species. At the cellular level, such as within endocrine cells or neurons, CREB can function as either a transcription activator or a transcription repressor. CREB dimerization and specific binding to the palindromic consensus cis-regulatory element, called CRE (cAMP responsive element; TGACGTCA), or to variant motifs present in the regulatory regions of target genes, are highly influenced by the surrounding DNA sequences and the extra-DNA environment (17–20). In addition, the specificity of CREB functionality is impacted by its own phosphorylation, acetylation, ubiquitination, sumoylation, and glycosylation (7, 8, 13, 16, 21–23). These and other regulatory mechanisms, such as those mediated by epigenetic elements and by non-coding small RNAs, facilitate CREB modulation of a wide variety of biological functions. These range from development to plasticity to disease, and include circadian rhythms, immune responses, and neuronal-related processes (14, 17, 18).

In rat, the multisynaptic circadian timing system (CTS) exerts its control over the PG via nighttime release of norepinephrine (NE) from sympathetic nerve endings stemming from neurons located in the superior cervical ganglia (SCG) (1, 2, 4). NE binds to β1 and α membrane, which triggers cooperative signaling cascades that end with the transient cAMP-induced phosphorylation of nuclear CREB1 at the Ser^133^ residue (pSer^133^-CREB1), as well as other related processes (4, 7, 8, 12, 22, 23). Once formed, pSer^133^-CREB1 initiates specific gene expression within the pinealocyte nuclei. In particular, pSer^133^-CREB1 induces *aa-nat* (arylalkylamine-N-acetyltransferase) gene expression that yields AA-NAT protein, which is one of the rate-limiting enzymes in melatonin synthesis (24–26). Additionally, thousands of other genes are regulated by NE-triggered cascades (27–31). These genes are involved, not only in hormone production, but also in many other pineal biological processes as well.

A new frame for understanding pineal biology was recently provided by sequencing the transcriptome of individual cells (28, 30). This study at single-cell resolution confirmed the high cellular heterogeneity within the rat PG, as it was previously proposed (32–36). These PG cell types were found to be transcriptionally distinct: two melatonin-producing pinealocyte subtypes (alpha and beta), three astrocyte subpopulations, two microglia subtypes, vascular and leptomeningeal cells, and endothelial cells (30). Alpha-pinealocytes are less abundant than beta-pinealocytes, but they synthesize nighttime melatonin more efficiently due to their higher capacity for O-methylating the precursor N-acetylserotonin (NAS). CREB1 transcripts were present in all nine cell types described above, but with different expression levels (30). In addition, no significant daily changes in *creb1* gene expression were found in any PG cell type. This is consistent with the reported observation that total CREB protein levels remain relatively constant throughout the light:dark (L:D) cycle in the whole rat PG (7). The reversible nocturnal phosphorylation of CREB1 at the Ser^133^ residue and pSer^133^-CREB1-dependent signaling pathways only provide a partial explanation for the highly specific behavior of this essential TF in the rat PG. This suggests that other regulatory mechanisms should be considered and studied. Our understanding is not yet clear about CREB1 spatio-temporal distribution and how CREB1 interacts with the chromatin and interchromatin elements of the pinealocyte nuclei. Dynamic spatial distribution of transcription factors is considered a feature of the highly plastic and compartmentalized structure of a cell nucleus, and it is expected to impact in nuclear functions (37–44).

Single-molecule studies of CREB binding and dissociation to its target sequence CRE, both *in vitro* and in living cells such as cortical neurons, have shown that CREB resides transiently and repetitively in fixed nuclear locations (hot spots) in the time range of several seconds (45, 46). This spatially restricted interaction takes place even though CREB acts as a mobile TF within the nuclear space (47). Kitagawa et al. also showed that the frequency of CREB binding to the highly localized genome spots was enhanced by neuronal activity, while CREB residence time on its nuclear target sites was unaffected (46).

The aim of our work presented herein was to study the spatio-temporal dynamics of nuclear CREB1 at single-cell resolution in the adult rat PG, and the nocturnal norepinephrine as a neural stimulus of this CREB1 behavior.

## Materials and methods

### Animals

All animal procedures performed in this study followed the U.S. National Institutes of Health’s Guide for Care and Use of Laboratory Animals and the Animal Research: Reporting *in vivo* Experiments (ARRIVE) Guidelines, and they were approved by the Institutional Animal Care and Use Committee (IACUC) at the School of Medicine, National University of Cuyo, Mendoza, Argentina (Protocol IDs 9/2012 and 74/2016). All efforts were made to minimize animal suffering. Three-month-old male Wistar rats were raised in our colony under controlled conditions, with 12:12 light:dark (L:D) cycle, and with *ad libitum* access to food and water. Room lights were turned on at 7 a.m. (*Zeitgeber* time 0; ZT0), and they were turned off at 7 p.m. (ZT12). Rats were euthanized by decapitation after ketamine/xylazine (50 and 5 mg/kg of body weight, respectively) anesthesia (48). Daytime pineal glands (PG), when no melatonin production was expected, were collected at ZT6 (middle of the light phase) and ZT10 (two hours before the lights were off). At night, samples were obtained under dim red light at ZT14 (early night) and ZT18 (middle of the dark phase), during the NE-induced transcription onset and high melatonin synthesis phases, respectively. Collected PG were parsed into groups for either Western blot (WB) analysis or for fluorescence immunohistochemistry (IHC).

### Surgical removal of superior cervical ganglia

Bilateral superior cervical ganglionectomy (SCGx) was performed on 3-month-old male Wistar rats, according to a previously described protocol (29, 32, 34, 48). Sham animals were subjected to surgery that included all the steps for exposing both superior cervical ganglia (SCG), but the actual excision was omitted. SCGx and sham-operated animals (N = 4 per each group) were housed in a stress-free environment for three weeks to prevent pinealocyte activation by stress-induced catecholamines. The Wallerian degeneration of the sympathetic nerve fibers from the SCG and the subsequent inflammatory environment within the PG were ameliorated at this post-surgical time point (34, 48). Following this surgical recovery, the rats were sacrificed at ZT14 (early night). The pineal glands from both groups were processed for fluorescence IHC.

### Western blot

For CREB1 detection by WB, total protein extracts from three independent pools of 10 PG each were generated at ZT6, ZT14 and ZT18 (N = 3 per each ZT). The rate-limiting enzyme in melatonin synthesis, the arylalkylamine-N-acetyltransferase (AA-NAT), was used to confirm the rhythm in melatonin production by the PG and simultaneously, the animal synchronization to the 12:12 L:D cycle (26). Actin was used as a loading control. To ensure protein identification, the extraction was performed using a two-step procedure in Triton X-100 Lysis Buffer [LB: 50 mM Tris-HCl, 150 mM NaCl, 5 mM EDTA, 0.5% (v/v) Triton X-100, pH 7.5], which was supplemented by adding phosphatase and protease inhibitors: 10 mM sodium fluoride (NaF), 10 mM sodium orthovanadate (Na3VO4), 1 mM phenylmethylsulfonyl fluoride (PMSF), and one protease inhibitor cocktail tablet (Cat# 11836153001, Roche Applied Science, Mannheim, Germany). Briefly, each PG pool was subjected to two successive treatments with cold LB to increase the protein yield. For the first treatment, the PG pool was homogenized in 150 microliters of cold LB at minimum speed, using a Bio-Gen PRO200 homogenizer (PRO Scientific Inc., Oxford CT, USA). The homogenate was centrifuged at 13,000 rpm, at 4°C for 15 minutes. The supernatant was collected and kept at 4°C as an initial protein concentrate. The remaining pellet was further subjected to a second 100-μL-cold LB treatment and then centrifuged at 4°C. The supernatants from both treatments were then combined to yield the final protein concentrate. The pellet resulting from the second treatment was discarded. Protein concentrations were estimated using the PierceTM BCA Protein Assay Kit (Cat# 23225, Pierce Biotechnology, Thermo Fisher Scientific Inc., Waltham, MA, USA). Proteins, at 50 micrograms per lane, were separated on sodium dodecyl sulfate (SDS)-polyacrylamide gels at 10% for CREB1, and at 12% for both AA-NAT and actin. Separated proteins were then transferred to PVDF membranes by semi-dry electroblotting. Membranes were incubated with blocking solution [5% (w/v) bovine serum albumin (BSA) in Tris-buffered saline (TBS) with 0.1% (v/v) Tween-20 (TBST) for CREB1, and 5% (w/v) skim milk in phosphate buffer saline (PBS) with 0.1% (v/v) Tween-20 (PBST) for both AA-NAT and actin]. A protein molecular weight marker was also loaded (PageRuler Plus Prestained Protein Ladder, Cat# 26619, Thermo Fisher Scientific Inc.). After rinsing, membranes were incubated overnight at 4°C with the following primary antibodies: rabbit monoclonal anti-CREB1 1:2,000 diluted in the corresponding blocking solution (5% BSA-TBST) (Cat# 9197, RRID: AB_331277, immunogen: recombinant protein specific to the amino terminus of human CREB1 protein, Cell Signaling Technology, Danvers, MA, USA); rabbit affinity isolated anti-actin 1:3,500 diluted in PBST (Cat# A2066, RRID: AB_476693, immunogen: synthetic peptide corresponding to the C-terminal actin fragment SGPSIVHRKCF, Sigma-Aldrich, St. Louis, MO, USA), and rabbit polyclonal anti-AA-NAT 1:15,000 diluted in PBST (AB3314, RRID: AB_2616598, immunogen: rat AA-NAT position 25-250; kindly provided by Dr. David C. Klein from NICHD, NIH, Bethesda, MD, USA). The secondary antiserum used was a donkey anti-rabbit antibody conjugated with horseradish peroxidase (HRP) (Cat# 711-035-152, RRID: AB_10015282, Jackson Immuno Research Labs, West Grove, PA, USA), dilution 1:50,000. Protein bands were visualized with the LAS-4000 system (ImageQuantTM LAS-4000, GE Healthcare Life Sciences, Pittsburgh, PA, USA) after a chemiluminescent reaction (Immobilon® Western Chemiluminescent HRP Substrate, Cat# WBKLS0100, EMB Millipore, Burlington, MA, USA). Total protein normalization was applied to compare levels of CREB1 and AA-NAT among ZTs, by using a modified procedure of the Coomassie blue staining method (49). Briefly, the blotted PVDF membranes were rinsed twice in TBS with 0.1% (v/v) Tween-20, and then stained for 1 minute with 0.1% (w/v) Coomassie brilliant blue R-250 (CBBR, Cat#1610400, Bio-Rad Laboratories Inc., Hercules, CA, USA) in methanol/Milli-Q water (1:1). The membranes were then successively destained for 2 minutes in acetic acid/ethanol/Milli-Q water (1:5:4), washed with Milli-Q water, and finally air-dried. The dry membranes were scanned with the LAS-4000 system. Densitometric analysis was carried out using the Image Lab SoftwareTM 6.0.1 (Bio-Rad Laboratories Inc.). The blots and the CBBR-stained membranes were merged *in silico*, and background was subtracted. To compensate for any differences in loading, the specific bands for CREB1 and AA-NAT that had been detected in the blots were normalized to the total CBBR-positive protein bands present in the corresponding lanes of the stained membranes. This normalization method is preferred over the older housekeeping protein procedure (50, 51).

### Fluorescence immunohistochemistry

Collected pineal glands (PG) were fixed by immersion in 4% paraformaldehyde (PFA) in PBS at 4°C and subsequently processed for immunostaining as described (32, 34, 35). Briefly, fixed PG were subjected to progressive dehydration, and then were embedded in Histoplast (Cat# 1203.59, Biopack, Bs. As., Argentina). Then, randomly oriented 10 μm-thick sections were cut from the middle region of each PG using a Microm HM 325 microtome (Thermo Fisher Scientific Inc.). For antigen retrieval, PG sections were boiled in 10 mM sodium citrate buffer (pH 6) containing 0.05% (v/v) Tween-20 for 30 min. Non-specific labeling was blocked by using 10% (v/v) donkey serum, 1% (v/v) Triton X-100 and 0.2% (w/v) gelatin in PBS, for 1 hour at room temperature (RT) in a humid chamber. After that, immunolabeling was performed by overnight incubation at RT in antibody buffer [2% (v/v) donkey serum, 1% (v/v) Triton X-100, and 0.2% (w/v) gelatin in PBS], which contained specific primary antibodies (Table 1).

**Table 1.**
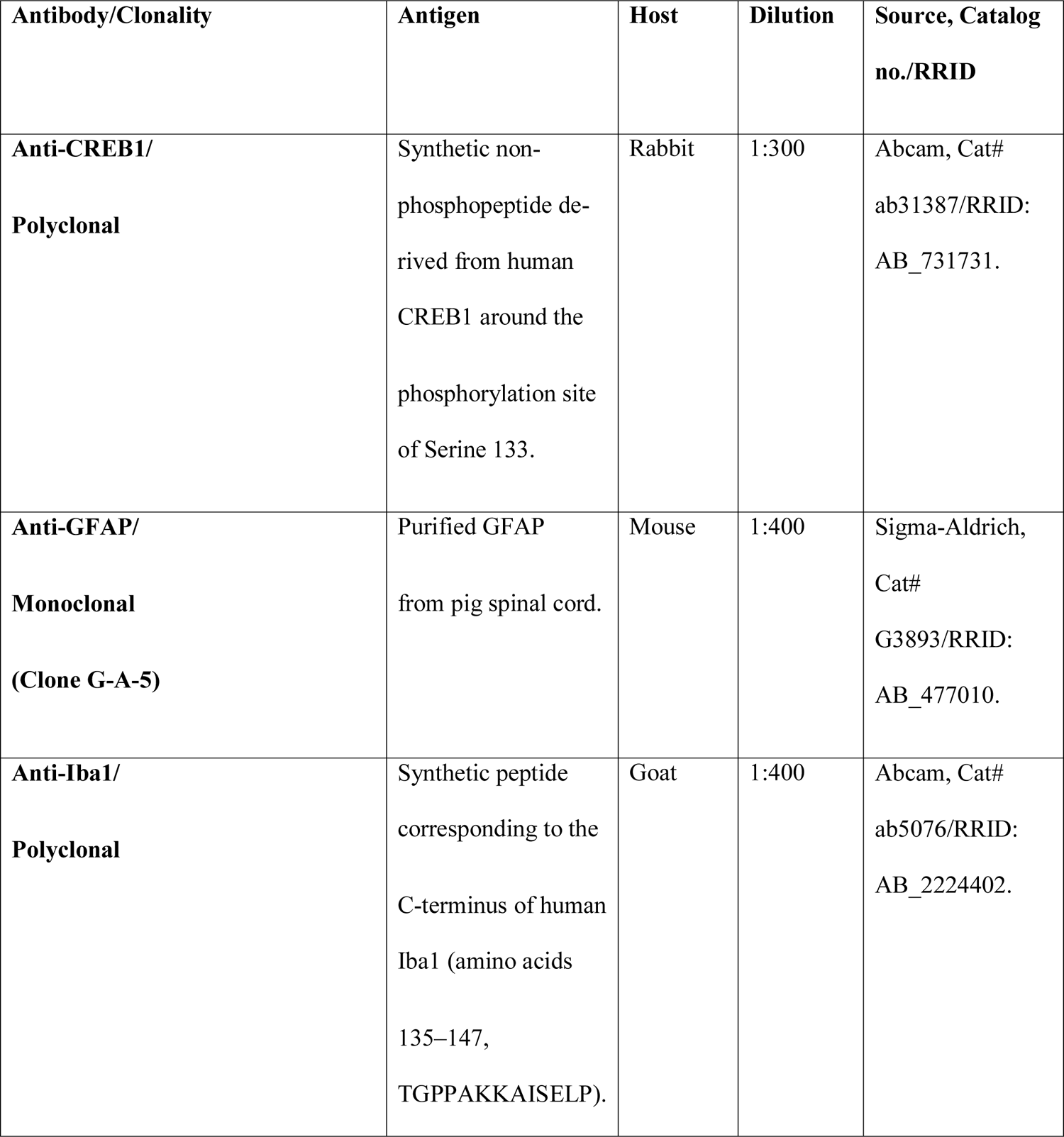
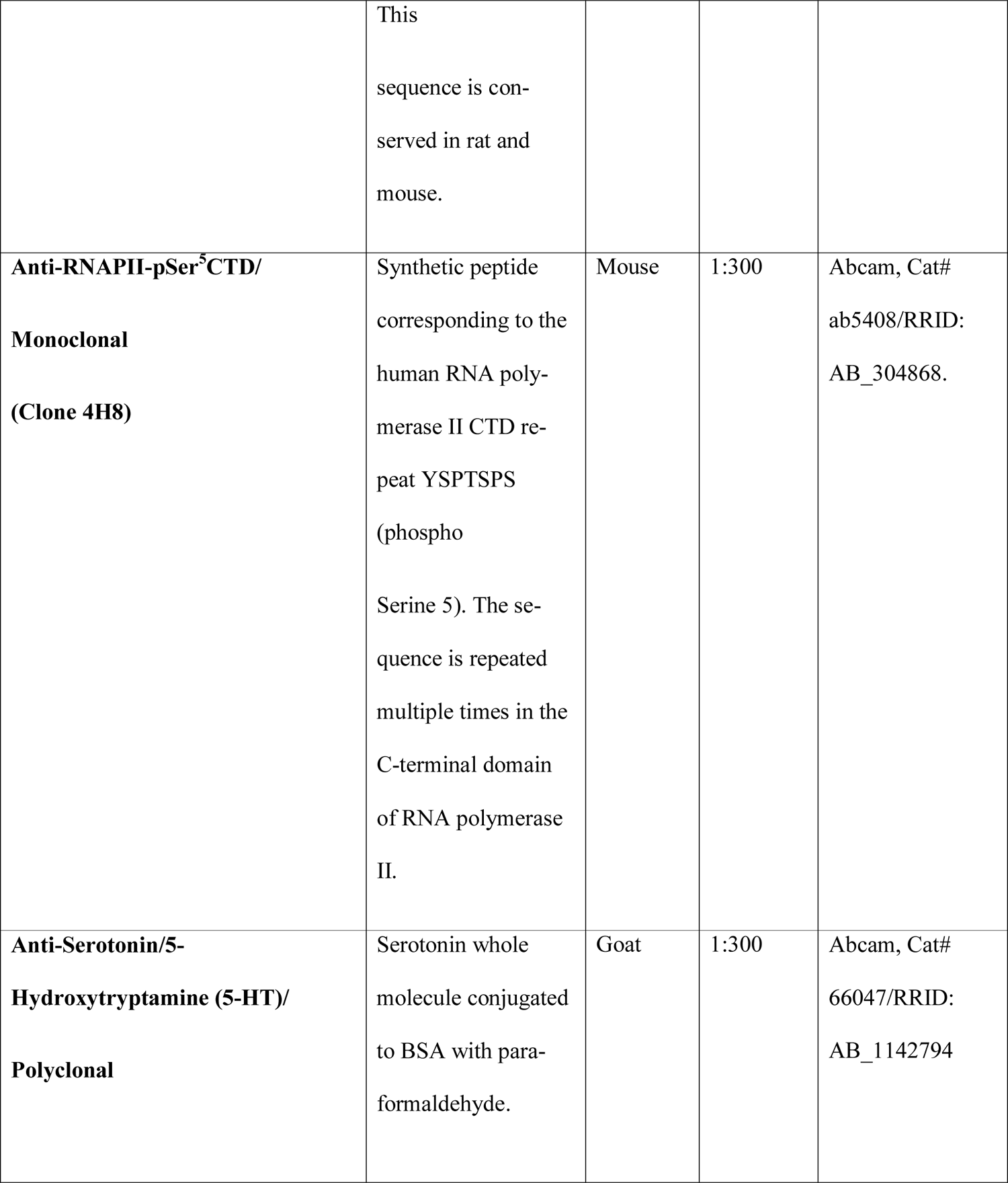
Primary antibodies for immunohistochemistry.

Following primary antibody incubation, sections were rinsed in PBS. Slices were then dipped in the secondary antibody-containing buffer with the nuclear dye 4′,6-diamidino-2-phenylindole (DAPI, Cat# D1306, Life Technologies-Invitrogen, Carlsbad, CA, USA, dilution 1:400), for 2 hours at RT. Secondary antibodies with low cross-reactivity, generated in donkey and conjugated with Alexa Fluor 488, Cy3 and Alexa Fluor 647, were used in different dilutions (Table 2).

**Table 2.**
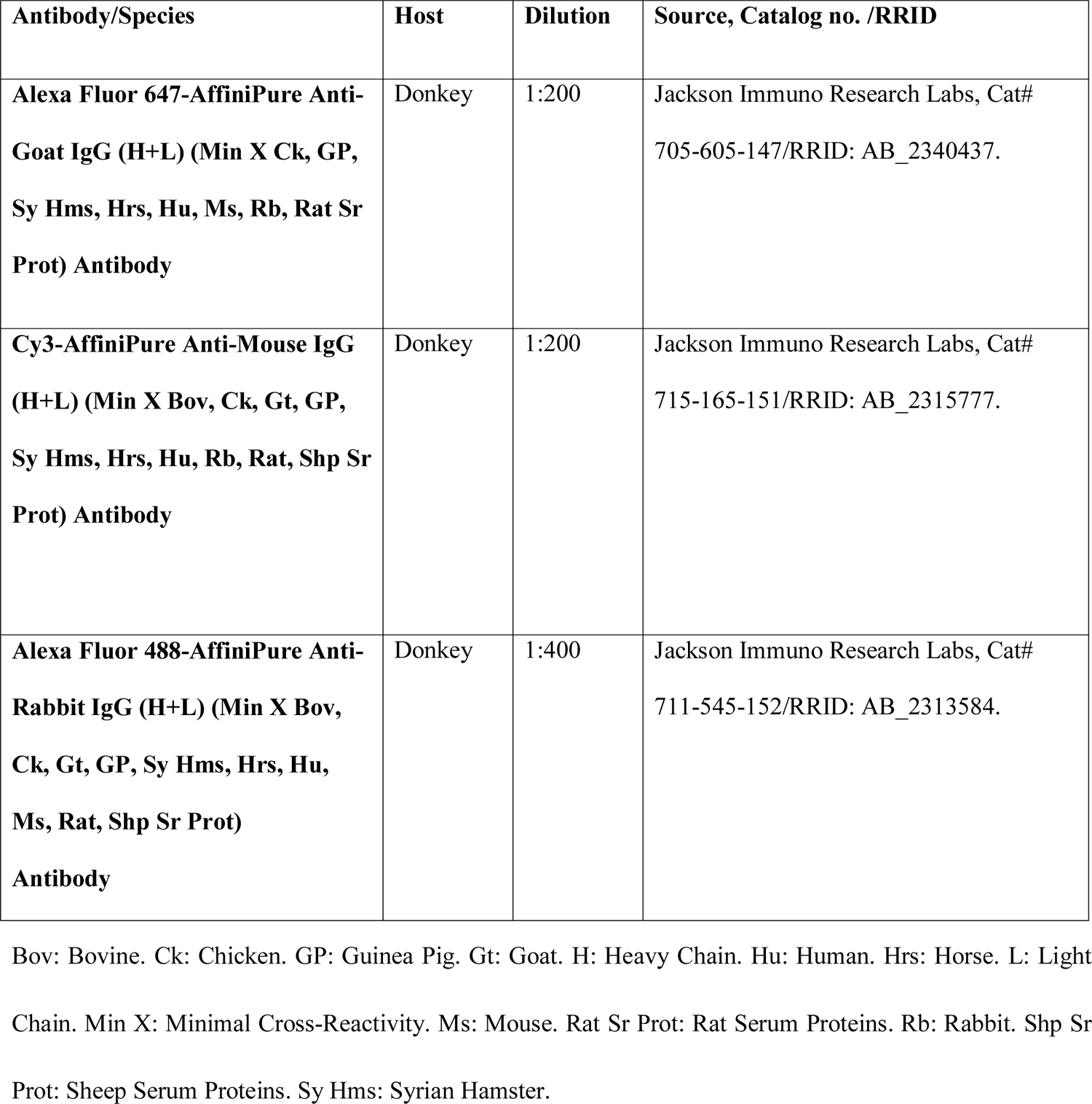
Secondary antibodies for immunohistochemistry.

| Antibody/Species | Host | Dilution | Source, Catalog no. /RRID |
| --- | --- | --- | --- |
| **Alexa Fluor 647-AffiniPure Anti-Goat IgG (H+L) (Min X Ck, GP, Sy Hms, Hrs, Hu, Ms, Rb, Rat Sr Prot) Antibody** | Donkey | 1:200 | Jackson Immuno Research Labs, Cat# 705-605-147/RRID: AB_2340437. |
| **Cy3-AffiniPure Anti-Mouse IgG (H+L) (Min X Bov, Ck, Gt, GP, Sy Hms, Hrs, Hu, Rb, Rat, Shp Sr Prot) Antibody** | Donkey | 1:200 | Jackson Immuno Research Labs, Cat# 715-165-151/RRID: AB_2315777. |
| **Alexa Fluor 488-AffiniPure Anti-Rabbit IgG (H+L) (Min X Bov, Ck, Gt, GP, Sy Hms, Hrs, Hu, Ms, Rat, Shp Sr Prot) Antibody** | Donkey | 1:400 | Jackson Immuno Research Labs, Cat# 711-545-152/RRID: AB_2313584. |

Bov: Bovine. Ck: Chicken. GP: Guinea Pig. Gt: Goat. H: Heavy Chain. Hu: Human. Hrs: Horse. L: Light Chain. Min X: Minimal Cross-Reactivity. Ms: Mouse. Rat Sr Prot: Rat Serum Proteins. Rb: Rabbit. Shp Sr Prot: Sheep Serum Proteins. Sy Hms: Syrian Hamster.

Sections were rinsed in PBS and covered with Mowiol mounting medium [9.6% (w/v) Mowiol 4-88 (Cat# 81381, Sigma-Aldrich) and 24% (v/v) glycerol in 0.1 M Tris–HCl buffer (pH 8.5)]. When the Cy3-conjugated secondary antibody was omitted, DAPI and Mowiol mounting medium were replaced by propidium iodide (PI, Cat# P4170, Sigma-Aldrich) and NPG-glycerol mounting medium [2% (w/v) N-propyl gallate (NPG, Cat# P3130, Sigma-Aldrich), 90% (v/v) glycerol, and 0.15% (w/v) propidium iodide in PBS], respectively. The controls of non-specific binding were routinely performed either by omitting primary antibodies, or by using blocking peptides when they were available (34, 35). Serial dilutions of each primary antibody alone were assayed to define the optimal antiserum concentrations. Double immunostaining was then performed, and the results were compared to those obtained from single antibody reactions. Imaging was done on an Olympus FV1000 (Olympus America Inc., Center Valley, PA, USA) confocal microscope. Images were processed with the ImageJ software (Version 1.52d, NIH, USA) and edited with Adobe Photoshop 7.0 (Adobe Systems Inc., San Jose, CA, USA).

### Analysis of CREB1 spatio-temporal dynamics within individual pinealocyte nuclei

To study CREB1 spatio-temporal dynamics within individual pinealocyte nuclei, PG sections were immunolabeled for CREB1 and then counterstained with DAPI. Images were acquired with an Olympus FV1000 confocal microscope, through a 60x/NA1.42/oil objective lens. A 2x digital zoom was applied during scanning to facilitate CREB1 analysis. Microscope parameters were set by sequential imaging of representative PG sections under non-saturated illumination conditions, with slices taken from control, sham-operated, and SCGx PG. For the detection of DAPI, a 405-nm laser at 4% intensity and 425 volts was used. This was followed by illumination with a 473-nm laser at 9% intensity and 570 volts to detect Alexa Fluor 488-labeled CREB1. The acquisition speed was 8 μ defining the scanning parameters, z-stack images were captured from three to four PG per ZT and per surgical condition (Table 3).

**Table 3.**
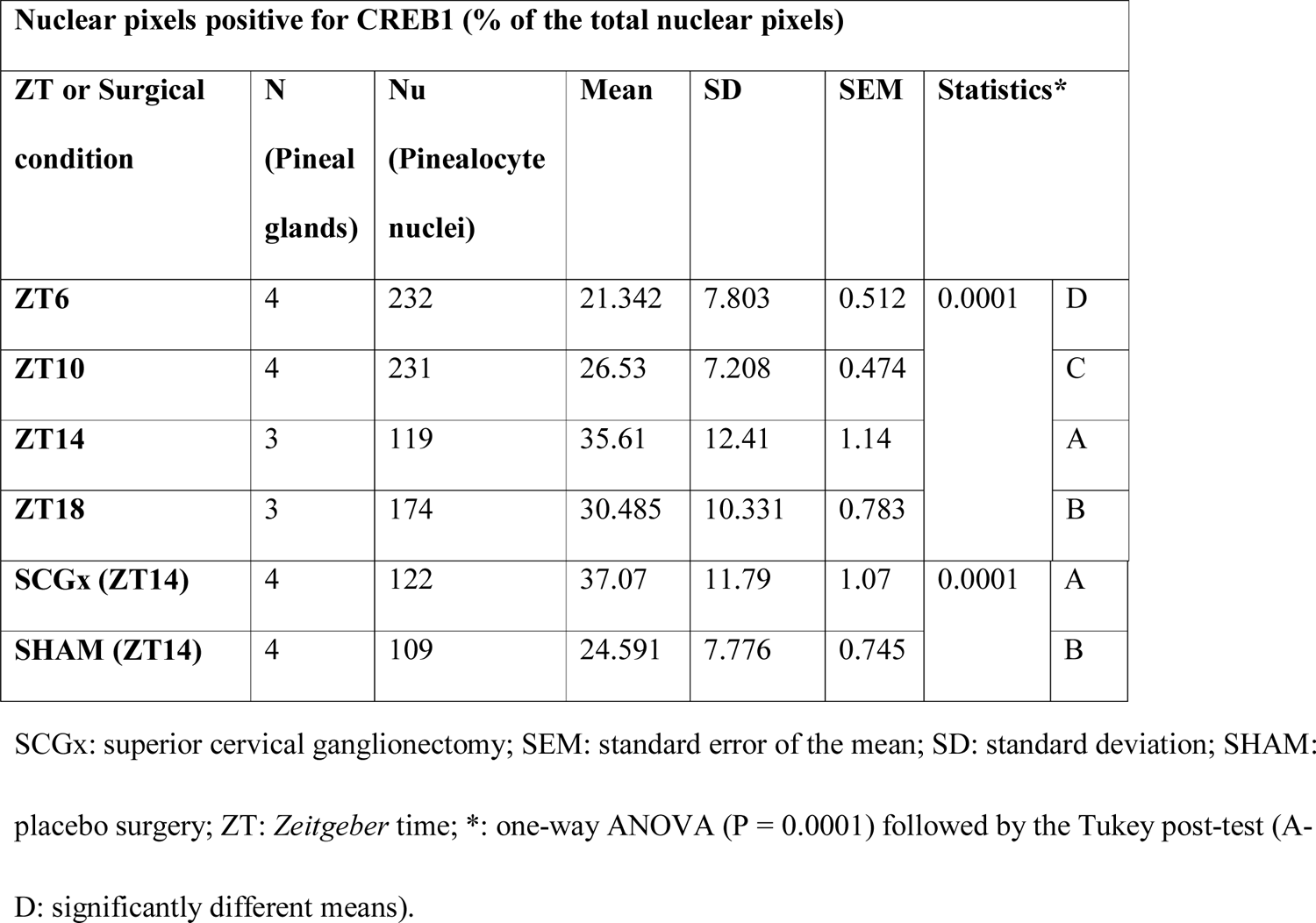
CREB1 distribution within individual pinealocyte nuclei

| Nuclear pixels positive for CREB1 (% of the total nuclear pixels) |  |  |  |  |  |  |  |
| --- | --- | --- | --- | --- | --- | --- | --- |
| ZT or Surgical condition | N (Pineal glands) | Nu (Pinealocyte nuclei) | Mean | SD | SEM | Statistics* |  |
| ZT6 | 4 | 232 | 21.342 | 7.803 | 0.512 | 0.0001 | D |
| ZT10 | 4 | 231 | 26.53 | 7.208 | 0.474 |  | C |
| ZT14 | 3 | 119 | 35.61 | 12.41 | 1.14 |  | A |
| ZT18 | 3 | 174 | 30.485 | 10.331 | 0.783 |  | B |
| SCGx (ZT14) | 4 | 122 | 37.07 | 11.79 | 1.07 | 0.0001 | A |
| SHAM (ZT14) | 4 | 109 | 24.591 | 7.776 | 0.745 |  | B |

SCGx: superior cervical ganglionectomy; SEM: standard error of the mean; SD: standard deviation; SHAM: placebo surgery; ZT: *Zeitgeber* time; \*: one-way ANOVA ( $P = 0.0001$ ) followed by the Tukey post-test (A-D: significantly different means).

For every PG section, four different areas were randomly chosen for z-stack image scanning. Z-stack images consisting of six focal planes in the z-axis with 1-μm steps, were collected as 1024×1024-pixel scans and then saved as .oib files (Olympus format). Pinealocytes were easily distinguished from interstitial cells because of their spatial organization in cords, as well as the characteristics of their nuclei including size, shape, euchromatin and heterochromatin ratio and distribution, and the presence of multiple nucleoli [32, 36]. On the other hand, the nuclei of most of the PG interstitial cells exhibited a highly homogeneous and compacted chromatin [35]. Identification of pinealocytes was also confirmed in adjacent sections within the same PG by immunostaining for serotonin or 5-hydroxytryptamine (5-HT), which is a melatonin precursor (S1 Fig) [35]. CREB1 fluorescence distribution and intensity in individual pinealocyte nuclei were determined in the z-stack images using the ImageJ software (Version 1.52d, NIH, USA). Briefly, the .oib files were transformed into .TIFF files, then stacked in the z-axis, then converted to 8-bit grayscale images, and finally they were subjected to the Otsu’s thresholding method to generate a binary mask per each channel (52). The resulting masks were merged with the original 8-bit z-stack images to extract image signal information and suppress any non-specific background. The corrected images were used for further characterization of separate pinealocyte nuclei. The ImageJ brush tool was applied to paint whole individual pinealocyte nuclei in the binary DAPI images and exclude those nuclei that were not properly individualized. The new masks were merged with the respective previously corrected 8-bit grayscale CREB1 and DAPI images, and the randomly selected individual pinealocyte nuclei were cropped to 200×200 pixels in size by using the ImageJ rectangle tool. In control animals, the total numbers of pinealocyte nuclei that were analyzed, were as follows: 232 at ZT6, 231 at ZT10, 119 at ZT14, and 174 at ZT18 (Table 3). In the SCGx and the sham-operated groups, the total numbers of pinealocyte nuclei were 122 and 109, respectively (Table 3). The ImageJ color histogram tool was used to determine the area for both CREB1 and DAPI within the cropped nuclei by counting the number of pixels occupied for each of these two species. Pixels occupied by CREB1 were normalized by expressing them as a percentage of the total nuclear pixels given by DAPI staining, and then were subjected to statistical analysis. CREB1 fluorescence intensity ranged from 0 to 255 for each pixel within the nucleus. These pixel intensity values were color look-up table (LUT) mapped and schematically represented using ImageJ interactive 3D surface plot tool.

### Statistical analysis

The minimal number of pinealocyte nuclei to be analyzed was defined for each ZT and for each surgical condition by using Minitab® 16.1.0 (Minitab® Statistical Software, State College, PA, USA). The normal distribution of the data was confirmed with the Anderson-Darling test (P > 0.01; Minitab® 16.1.0). Data, as expressed by the mean ± SEM (standard error of the mean), were analyzed using PRISM 6 (GraphPad Software Inc., La Jolla, CA, USA) (Table 3). One-way ANOVA followed by the Tukey post-test was performed. A P < 0.05 was considered significant.

## Results

First, we examined the abundance of total CREB1 in the rat pineal gland by Western blot (WB) during the light phase (ZT6) and the dark phase (ZT14 and ZT18) of a 12:12 L:D cycle (Fig 1). Our results confirmed previous reports that showed that total CREB protein does not manifest a daily rhythm in the rat PG (one-way ANOVA; P = 0.83) (Fig 1B) [7]. The rhythmic nature of the rat PG was verified by detection of the nocturnal AA-NAT (one-way ANOVA; P = 0.0008) (Fig 1B), which is one of the rate-limiting enzymes in the melatonin synthesis [26].

**Figure 1.**
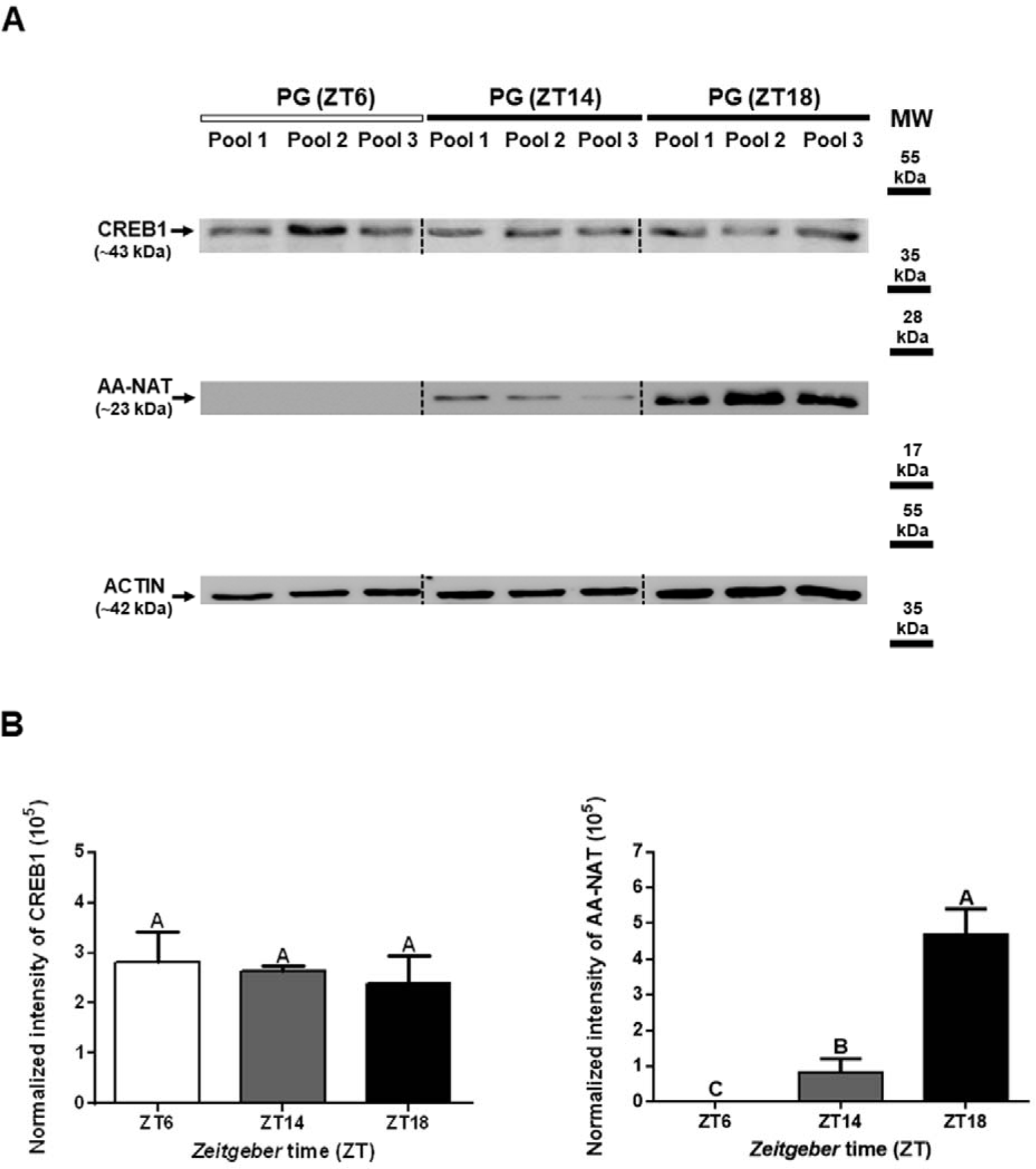
CREB1 levels in the rat pineal gland during the daytime and the nighttime. Pools of 10 adult rat pineal glands (PG) were collected during each of three different *Zeitgeber* times: ZT6 (middle of the light phase), ZT14 (early night), and ZT18 (middle of the dark phase). A total protein extract was prepared from each pool for Western blot (WB) analysis of the following: the cyclic AMP (cAMP) responsive element-binding protein 1 (CREB1), the arylalkylamine-N-acetyltransferase (AA-NAT), and actin. AA-NAT is one of the rate-limiting enzymes in melatonin synthesis. For each ZT, three pools were analyzed, with 50 µg protein loading per lane. (A) Representative blots showing specific bands for CREB1 (∼43 kDa), AA-NAT (∼23 kDa), and actin (∼42 kDa). As expected, AA-NAT was mainly present in nighttime samples, and its expression increased during the dark phase. (B) The abundance of CREB1 and AA-NAT were normalized to the total amount of protein per lane, by using a modified procedure of the Coomassie blue staining method. Data were expressed as mean ± standard error of the mean (SEM). Statistics: one-way ANOVA (P = 0.83 for CREB1 and P = 0.0008 for AA-NAT) followed by the Tukey post-test (A: no significant differences were found among ZTs for CREB1; A-C: significantly different means for AA-NAT). MW: molecular weight.

Then, we aimed to study the spatio-temporal dynamics of CREB1 at the level of individual pinealocyte nuclei, using fluorescence immunohistochemistry followed by confocal laser-scanning microscopy and quantitative analysis. The highly specific ab31387 antibody against total CREB1 (53–55), showed that CREB1 was present in the nuclear compartment of all PG cell types. No fluorescent signal was observed in the cytoplasm of any of these cells under the non-saturated illumination conditions that were applied during image scanning. Despite the presence of nuclear CREB1 in all PG cells, pinealocytes emerged as having discrete CREB1 domains within their nucleoplasm that were especially distinct. The number, size, and location of the CREB1-immunoreactive nuclear foci varied among pinealocytes within the same PG (S2 Fig). In non-pinealocyte cells the distribution of CREB1 was more homogenous and denser. Interstitial cells immunoreactive for CREB1 were identified as phagocytes positive for microglia/macrophage-specific ionized calcium-binding adapter molecule 1 (Iba1) (S3 Fig), and as astrocyte-like cells enriched in glial fibrillary acidic protein (GFAP) (S4 Fig). The CREB1/GFAP double-positive cells were observed in the proximal pole of the PG, near the pineal stalk. Cells with elongated nuclei such as endothelial cells and fibroblasts, also exhibited a condensed distribution of CREB1 in their nuclei (S3 Fig).

Although pinealocyte nuclei exhibited a heterogeneous distribution pattern of CREB1 among them, daily variations were observed when daytime and nighttime samples were compared (Fig 2 and S5 Fig). Analysis of high-magnification z-stack images of individual pinealocyte nuclei revealed that CREB1-immunoreactive foci were significantly more dispersed in the dark phase (ZT14 and ZT18) than in the light phase (ZT6 and ZT10) (Figs 2 and 3, and S6-S9 Figs). For this analysis, a multi-step image processing method was applied using the ImageJ software (Version 1.52d, NIH, USA). The number of pixels occupied by CREB1 in a selective nucleus was normalized to the total nuclear pixels given by DAPI staining, and then it was expressed as a percentage. Pinealocyte nuclei with low, medium, and high percentages of CREB1-immunoreactive nuclear pixels were found in each ZT, consistent with the heterogeneity in CREB1 spatial distribution observed among pinealocytes (Fig 3). Interestingly, statistics confirmed the larger dispersion of CREB1 within individual pinealocyte nuclei at night (one-way ANOVA; P = 0.0001) (Fig 3E, Table 3). Quantitative analysis was not applied to non-pinealocyte cells due to the compact distribution of the fluorescent pixels for CREB1 in their nuclei, and the lack of observed daily variations for them.

**Fig 2.**
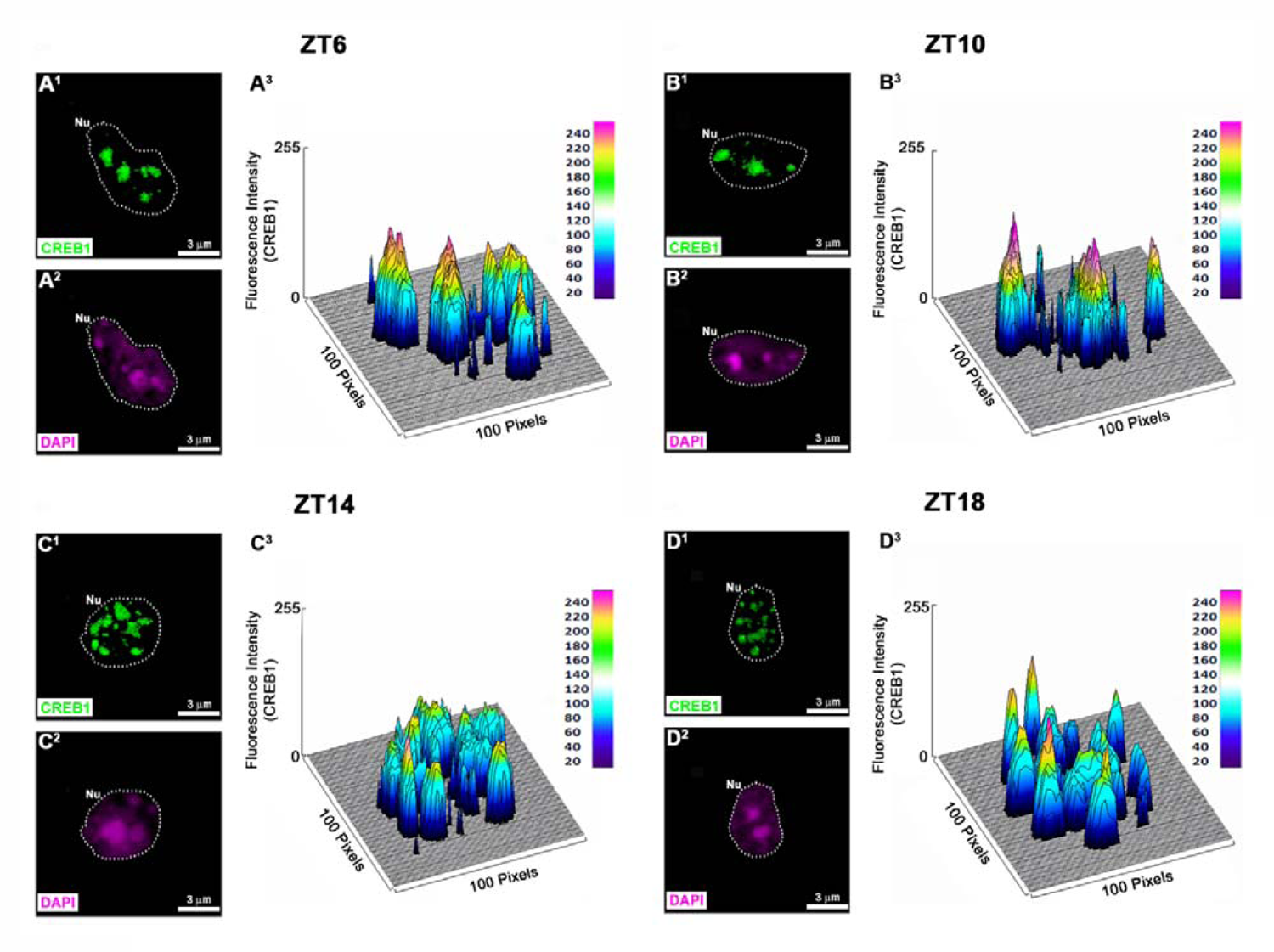
Heterogenous spatio-temporal distribution of CREB1 within the nuclei of rat pinealocytes. Representative separate pinealocyte nuclei (Nu) immunolabeled for CREB1 (green) and counterstained with 4′,6-diamidino-2-phenylindole (DAPI; magenta) are shown at daytime (ZT6 and ZT10), and at nighttime (ZT14 and ZT18). The nuclear perimeter is defined by a dashed white line. (A^1^, B^1^, C^1^, D^1^) Pinealocyte nuclei immunostained for CREB1. (A^2^, B^2^, C^2^, D^2^) The same nuclei shown in A^1^, B^1^, C^1^ and D^1^, but dyed with DAPI. (A^3^, B^3^, C^3^, D^3^) Schematic representations illustrating the fluorescence intensity of CREB1 for each pixel within the pinealocyte nucleus. The fluorescence intensity ranges from 0 to 255. The interactive 3D surface plot tool, included in the ImageJ software (Version 1.52d, NIH, USA), was used to build these reconstructions. (A^1^-A^2^, B^1^-B^2^, C^1^-C^2^, D^1^-D^2^) 2x digital zooms from 60x images; scale bar: 3 μm. ZT: *Zeitgeber* time.

**Fig 3.**
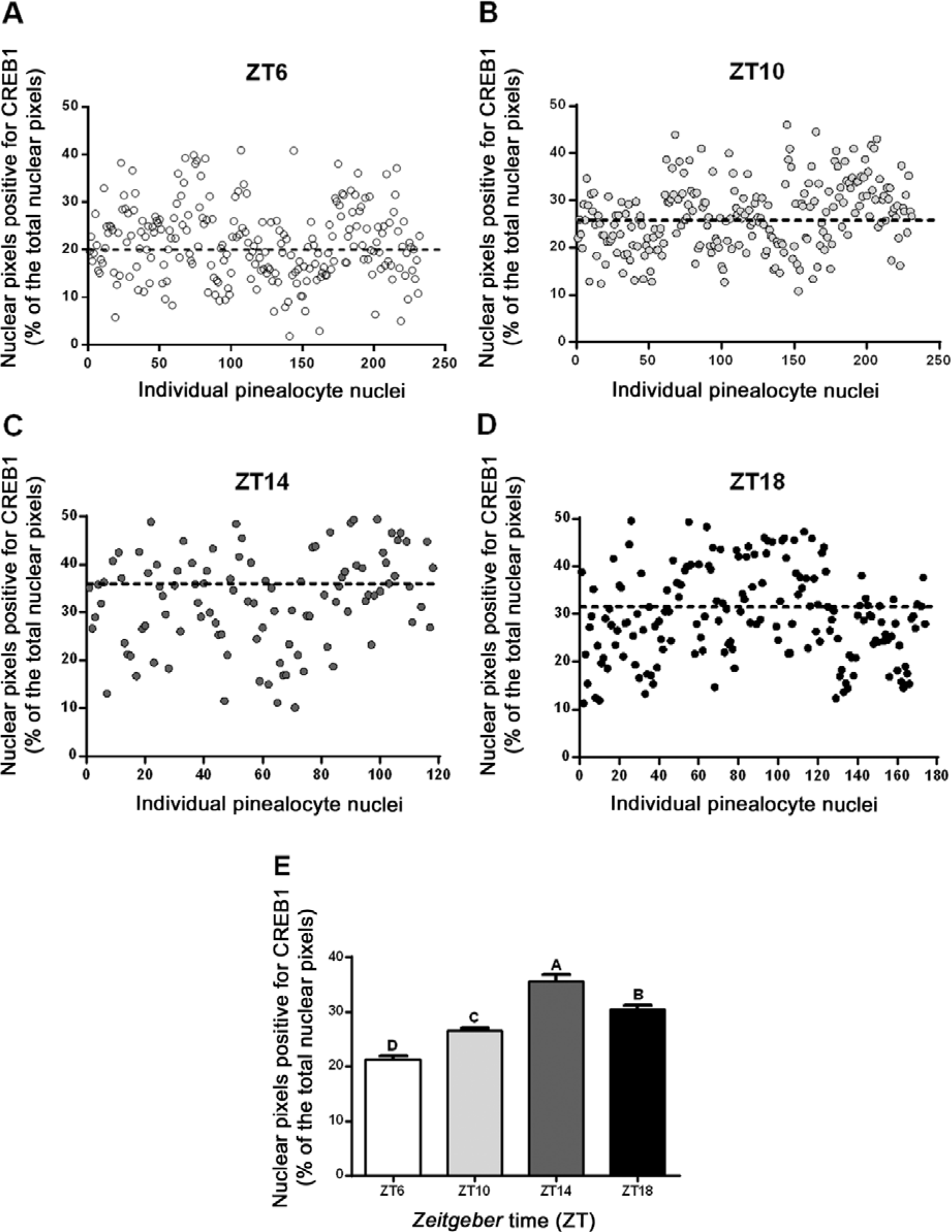
Daily rhythm in the nuclear distribution of CREB1 in rat pinealocytes. Pineal gland sections immunolabeled for CREB1 and counterstained with 4′,6-diamidino-2-phenylindole (DAPI) were imaged with a confocal microscope. Images were processed with the ImageJ software (Version 1.52d, NIH, USA) to identify and analyze separate pinealocyte nuclei. Three or four pineal glands per ZT were used. Total number of pinealocyte nuclei was: 232 at ZT6 (A), 231 at ZT10 (B), 119 at ZT14 (C), and 174 at ZT18 (D). Pixels occupied by CREB1 were quantified and were then normalized by expressing them as a percentage of the total nuclear pixels. (A-D) Each circle represents the percentage of pixels immunoreactive for CREB1 within an individual nucleus, at a defined ZT. The mean value for each ZT is represented by a horizontal dashed black line. High heterogeneity among pinealocyte nuclei is observed at each ZT. (E) Mean ± standard error of the mean (SEM). Statistics: one-way ANOVA (P = 0.0001) followed by the Tukey post-test (A-D: significantly different means). ZT: *Zeitgeber* time.

To study the potential link between the daily rhythm in CREB1 spatial distribution and the overall transcription activity within individual pinealocyte nuclei, the expression of a particular phosphorylated form of the RNA polymerase II (RNAPII) was studied by using a monoclonal antibody suitable for IHC. This antibody was raised against a synthetic peptide corresponding to the RNAPII C-terminal repeat domain (CTD) YSPTSPS, modified at the Ser^5^ residue (pSer^5^) (S10 Fig). The specific RNAPII-Ser^5^CTD signal was found with a wide and punctate distribution in the pinealocyte nucleoplasm, with the exception that no signal was detected in the nucleolar domains under the non-saturated illumination conditions that were applied during image scanning. No apparent differences in pinealocyte RNAPII-pSer^5^CTD expression were observed between ZTs. On the other hand, the RNAPII-pSer5CTD levels did vary among nuclei of non-pinealocyte cells within the same PG.

To determine if the nocturnal pulse of norepinephrine (NE) from the sympathetic nerve endings influences the spatial organization of CREB1 within individual pinealocyte nuclei, the transcription factor was studied at ZT14 (early night) in PG extracted from rats that were subjected to either chronic bilateral superior cervical ganglionectomy (SCGx) or placebo surgery (SHAM). High-resolution z-stack images of individual pinealocyte nuclei from SCGx and SHAM PG were processed and analyzed using the ImageJ software (Fig 4, and S11 and S12 Figs). Discrete CREB1-immunolabeled nuclear domains were observed in pinealocyte nuclei under both surgical conditions. As was found in the non-operated animals (Fig 3), both SCGx and SHAM states exhibited heterogeneity in the percentage of nuclear pixels immunoreactive for CREB1 among different pinealocytes within the same PG (Fig 4C and 4D). In the SCGx pinealocytes, however, nuclear CREB1 appeared significantly more dispersed, as compared to SHAM pinealocyte nuclei (one-way ANOVA; P = 0.0001) (Fig 4E, Table 3). In addition, at ZT14, the average percentage of nuclear pixels occupied by CREB1 did vary significantly between non-operated (control) and sham-operated rats (Fig 4E). No apparent differences in CREB1 nuclear distribution were observed in interstitial cells following the surgical procedures.

**Fig 4.**
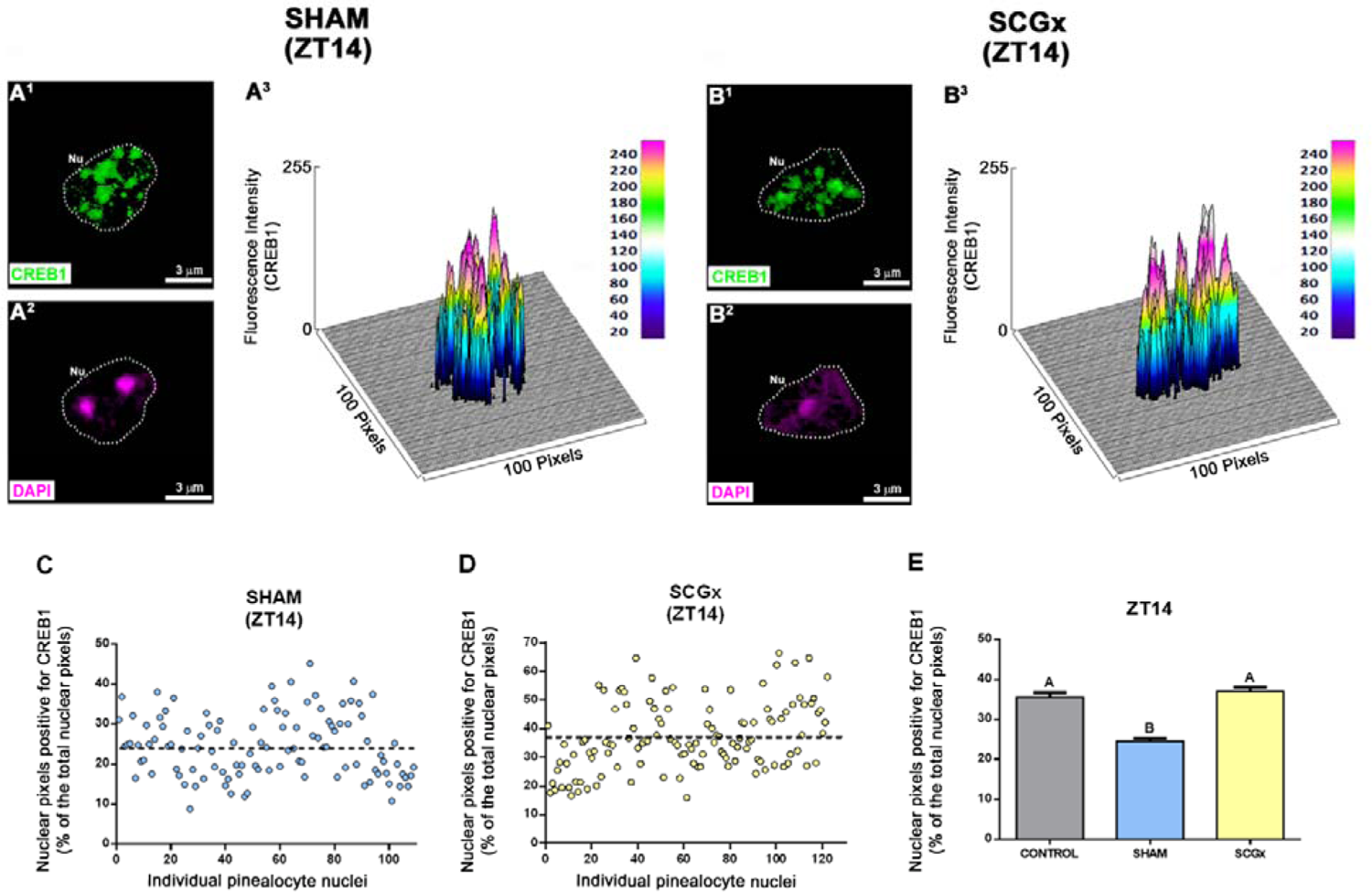
Spatial distribution of CREB1 within rat pinealocyte nuclei after chronic bilateral superior cervical ganglionectomy. Adult rats were subjected to either bilateral superior cervical ganglionectomy (SCGx; N=4) or placebo surgery (SHAM; N=4). All pineal glands (PG) were collected three weeks after surgeries at ZT14, and were then sectioned and immunolabeled for CREB1 (green). 4′,6-diamidino-2-phenylindole (DAPI; magenta) was used as nuclear dye. (A^1^, B^1^) Representative separate pinealocyte nuclei (Nu), positive for CREB1. (A^2^, B^2^) The same nuclei shown in A1 and B1, but stained with DAPI. The nuclear perimeter is defined by a dashed white line. (A^3^, B^3^) Schematic representations of the fluorescence intensity of CREB1 for each pixel within the pinealocyte nucleus. The fluorescence intensity ranges from 0 to 255. (A^1^-A^2^, B^1^-B^2^) 2x digital zooms from 60x images; scale bar: 3 μm. (C-E) Individual pinealocyte nuclei were identified and analyzed from confocal images using the ImageJ software (Version 1.52d, NIH, USA). Total numbers of pinealocyte nuclei were 122 for the SCGx group, and 109 for the SHAM group. Pixels occupied by CREB1 were quantified and were then normalized by expressing them as a percentage of the total nuclear pixels. (C, D) Each circle represents the percentage of pixels immunoreactive for CREB1 within an individual nucleus, under the defined condition. The mean value for each group is represented by a horizontal dashed black line. High heterogeneity among pinealocyte nuclei is observed in both groups. (E) Mean ± standard error of the mean (SEM). The control group included pineal glands from adult rats housed under a 12:12 light:dark (L:D) cycle, that were sacrificed at ZT14 (See Fig 3C). Statistics: one-way ANOVA (P = 0.0001) followed by the Tukey post-test (A, B: significantly different means). ZT: *Zeitgeber* time.

## Discussion

In this study, we describe for the first time the spatio-temporal dynamics of the transcription factor (TF) cyclic AMP responsive element-binding protein 1 (CREB1) at single-cell resolution within the mature rat pineal gland (PG). Two highly specific anti-CREB1 antibodies were used. These antibodies were raised against human CREB1 regions, which are highly homologous to rat CREB1 and its isoforms.

CREB is a widespread TF, yet its functionality differs from one cell type to another (56, 57). Nighttime neural stimulus-induced phosphorylation of nuclear CREB1 at one defined serine residue, Ser^133^, and the subsequent pSer^133^-CREB1 binding to CRE sites in the regulatory regions of key target genes, only partially explain the regulatory complexity behind the rhythmicity of pineal biology (7, 8, 12, 21–23). In fact, no daily changes in total CREB1 protein were found in the rat PG (Fig 1), as has been previously reported (7). In addition, no significant day/night differences were observed in CREB1 mRNA levels in any of the nine transcriptionally distinct PG cell types recently identified by Mays et al. (30). Taken together, the study of CREB1 nuclear distribution and CREB1 interaction with chromatin and interchromatin elements may represent a novel approach for further understanding the regulatory mechanisms behind PG rhythmicity. This approach might also be suitable for further analysis of other members of the CREB/CREM/ATF-1 family, including the well-characterized pineal repressor ICER and the under-studied CREB2, CREB3 and CREB3L2, and other TF families that are involved in pineal biology (9, 31, 58, 59).

Our analysis of total CREB1 spatial distribution revealed that CREB1 is present in the nuclear compartment of both the melatonin-producing pinealocytes and the non-pinealocyte cells of the adult rat PG (Fig 2, and S2-S5 Figs). In addition, no cytoplasmic CREB1 was detected for any PG cell type, under the non-saturated image scanning conditions that were used. Nevertheless, pinealocytes emerged as a distinct cell population due to CREB1 presence in restricted nuclear domains. This pattern was characteristic of CREB1. In contrast, for example, a homogeneous arrangement was observed for both transcription factors, the pinealocyte lineage-determining Pax6, and the ontogenetic and homeostatic NeuroD1 (32, 35). This suggests that the CREB1 pattern within pinealocyte nuclei may not be exclusively determined by the fact that the DNA itself is spatially heterogenous (S1 Fig).

Discrete nuclear hot spots immunoreactive for CREB were previously found and described in other neuronal cell types, such us mouse neuroblastoma Neuro2a cells and cortical neurons from 16-day-old mouse embryos (45, 46). These single-molecule studies have shown that CREB residency on the regulatory regions of its target genes is dynamic, and that CREB dwells there for a short duration, in the range of several seconds. Kitagawa et al. also showed that neuronal activity promoted CREB-dependent transcription by potentiating the frequency of CREB binding to well defined and highly specific genome locations (46).

Heterogeneity was observed in the number, size, shape, and location of the CREB1-immunoreactive nuclear foci among mature pinealocytes from the same PG (S2 Fig). On the other hand, all non-pinealocyte cells within the rat PG exhibited high levels of CREB1, that were uniformly distributed within their nucleoplasm. This CREB1 pattern was consistent with a homogenous and dense chromatin in these cell types. Higher microscope resolution imaging is needed, however, to better study CREB1 dynamism in these interstitial cells (39, 60, 61). Identified non-pinealocyte cells included Iba1-immunoreactive phagocytes, GFAP-positive astrocytes, as well as interstitial cells with elongated nuclei such as endothelial cells and fibroblasts (S3 and S4 Figs). These observed differences among PG cell types prompted us to focus solely on the pinealocyte population for quantitative analysis of CREB1 spatio-temporal dynamics. Therefore, a multi-step image processing method was developed using the ImageJ software, which uncovered dispersion in the percentage of nuclear pixels occupied by CREB1 within pinealocyte nuclei in the same PG. High, medium, and low values in the occupancy of nuclear pixels by CREB1 were observed for each ZT (Fig 3, and S6-S9 Figs). Statistically significant differences emerged, however, between the light and dark phases (Fig 3E, Table 3). In fact, a larger dispersion of the TF was observed at nighttime when the pineal environment is under the influence of neural norepinephrine (NE).

Binding and dissociation of transcriptions factors to their target genes and the impact on transcription itself, are certainly influenced by the surrounding chromatin. To better understand how pinealocyte CREB1 spatial distribution is linked to transcription, we studied the day/night expression of a particular phosphorylated form of the RNA polymerase II (RNAPII-pSer^5^CTD) which is a marker of the overall transcription activity. Changes in the RNAPII C-terminal repeat domain (CTD) code, such as phosphorylation and dephosphorylation, determine the assembly of different sets of nuclear factors and ultimately influence the functional organization of the nucleus (62, 63). Our results showed that RNAPII-pSer^5^CTD is widely expressed in the nucleoplasm of pinealocytes during both daytime and nighttime (S10 Fig). This is consistent with upregulated PG genes reported in both the light phase and the dark phase of the L:D cycle in rat (27–31). Using conventional confocal microscopy to examine pinealocytes, we found no relationship between the total RNAPII-pSer^5^CTD distribution pattern and the daily variations in CREB1-immunoreactive nuclear foci. However, higher resolution characterization of the discrete CREB1 nuclear domains will likely provide further information about the crosstalk between this TF and chromatin and interchromatin elements (39, 45, 46, 60, 61). In fact, it was previously proposed that transient NE-induced phosphorylation of one specific serine residue, Ser^10^, of histone H3, may facilitate nocturnal actions of pSer^133^-CREB and other TF in the rat PG (64, 65). Furthermore, acetylation of Lys^14^ (lysine 14) in histone H3 was found to have both inhibitory and stimulatory effects on NE-induced genes, adding even more complexity to PG gene expression regulation (66, 67). On the other hand, the mouse liver has been reported to have a robust peripheral circadian clock, with a stereotypical time-dependent pattern of transcriptional architecture and chromatin landscape (68, 69). These genome-wide analyses of the mammalian clock transcriptional feedback loops have revealed a global circadian regulation of transcription- and chromatin-related processes. Because the PG is a fundamental element of the circadian timing circuit, it might serve as an interesting model for studying the spatio-temporal dynamics and interrelationships of chromatin and interchromatin components, with transcription factors expected to be mainly found in the interchromatin compartment (37).

CREB is known to be a TF that is influenced by a wide variety of stimuli, including neurotransmitters related to neuronal activity (14, 16, 17, 46). In this PG study, we investigated the impact of nocturnal neural norepinephrine (NE) on the nuclear distribution of CREB1 in pinealocytes at the early night (ZT14). Adult rats were subjected to either bilateral superior cervical ganglionectomy (SCGx) or sham surgeries. Three weeks after these surgical procedures, pineal glands were collected and the spatial distribution of nuclear CREB1 was quantitatively analyzed (Fig 4, and S11 and S12 Figs). The percentage of nuclear pixels occupied by CREB1 did vary among pinealocytes from the same PG, under both surgical conditions (Table 3). However, CREB1 appeared more dispersed in pinealocyte nuclei after suppression of the nocturnal NE, as compared to sham-derived cells. In addition, differences in CREB1 distribution were found between sham-operated and non-operated (control) rats (Fig 4E). We speculated that these differences may stem from the surgical steps themselves or from the post-surgical recovery, as performed on the sham-operated rats. These results do not rule out the influence of NE on nuclear CREB1 dynamics, but they do add to the regulatory complexity of the PG transcriptional architecture. CREB1 analysis was applied when the Wallerian degeneration of the sympathetic nerve fibers from the SCG and the subsequent inflammatory environment within the PG were ameliorated, and the SCGx-induced microgliosis was resolved (34, 48). However, chronic SCGx might cause lasting milieu changes that could affect the crosstalk between pinealocytes and local phagocytes and might also affect the pinealocyte nuclear architecture and function. For a related example, an inflammatory insult caused by bacterial lipopolysaccharide (LPS) was shown to activate the microglia-pinealocyte network and to inhibit *de novo* gene expression in the pinealocyte nuclei, which in turn dampened melatonin synthesis (70). LPS-induced inflammation may also cause changes in nuclear CREB1 spatio-temporal dynamics and investigating this may shed further light about how cytokines impact the pinealocyte transcriptional architecture and their ultimate influence on nuclear functionality. Similarly, other stimuli, such as SCG decentralization and the administration of adrenergic agonists and antagonists, could also possibly affect CREB1 nuclear distribution and function.

The results presented in this study were summarized in the schematic representation of Figure 5 (Fig 5). The figure shows heterogeneity in the nuclear distribution of CREB1 protein among the different PG cell types. A daily dynamism of the nuclear domains occupied by CREB1 is clearly observed only in pinealocytes, with larger dispersion during the dark phase when the PG milieu is under the influence of neural norepinephrine (NE). On the other hand, chronic sympathetic disruption caused by bilateral SCGx, disperses CREB1 in the nocturnal pinealocyte nuclei, as compared to the sham-derived cells. Highly dense distribution of nuclear CREB1 in non-pinealocyte cells was not obviously affected by temporal cues, nor by the surgical procedures.

**Fig 5.**
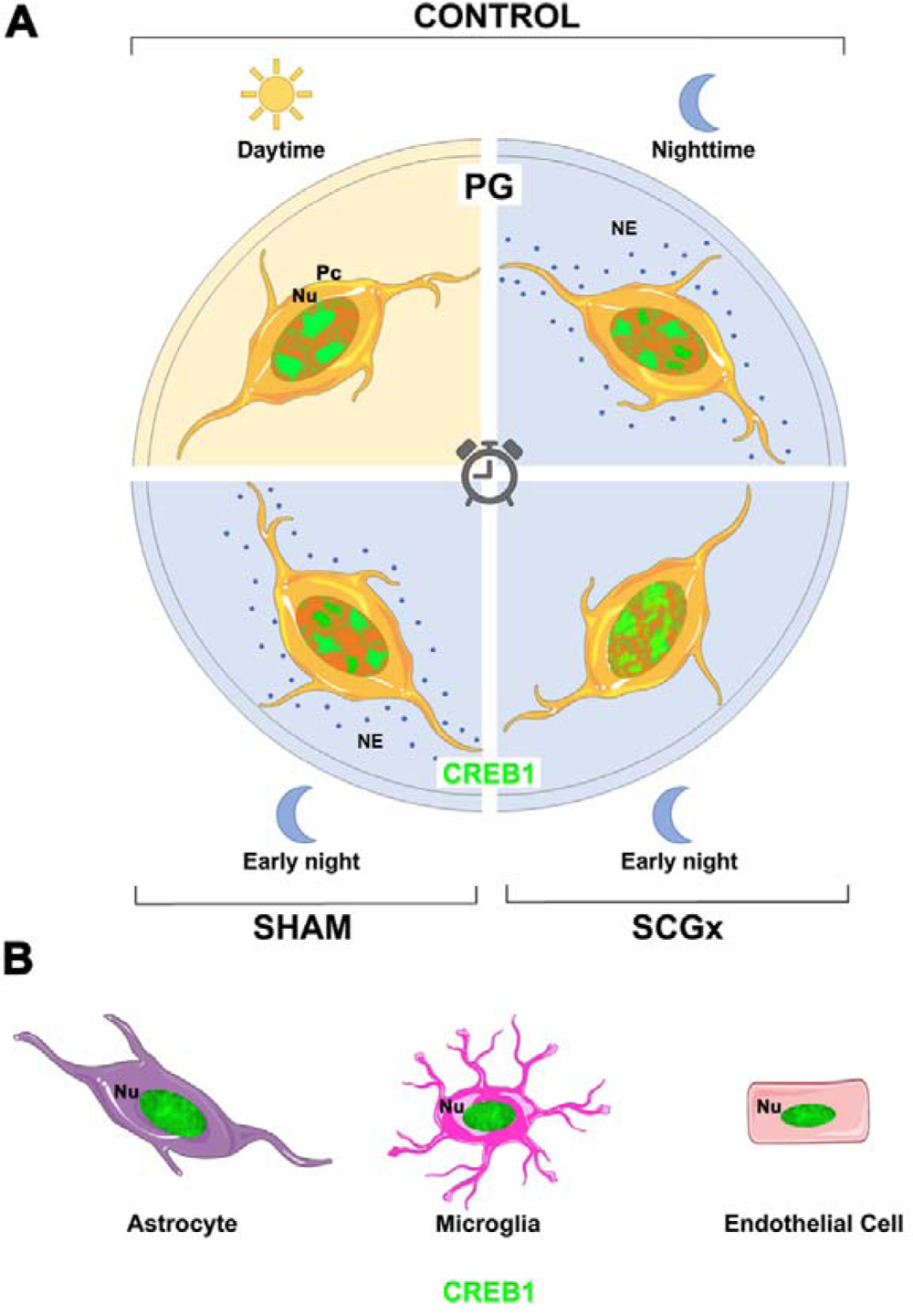
Schematic representation of the spatio-temporal dynamics of nuclear CREB1 within the rat pineal gland. The figure shows heterogeneity in the nuclear distribution of CREB1 protein (green) among the different pineal gland cell types, including pinealocytes (Pc, orange) and the following interstitial cells: astrocytes (violet), phagocytes (magenta), and cells with elongated nuclei such as endothelial cells (pink). (A) In the upper half of the model, pinealocytes identified in pineal glands (PG) from control rats housed under a 12:12 light:dark (L:D) cycle, are schematized at both daytime and nighttime. In the lower half of the model, pinealocytes are shown which were identified in PG from sham-operated (SHAM) and ganglionectomized (SCGx) animals, sacrificed three weeks after surgeries at early night (ZT14). Nuclear CREB1 exhibits a distinctive distribution pattern in pinealocytes. The spatial dynamism of CREB1 during the L:D cycle is clearly observed in the melatonin-producing cells. In the light phase, the transcription factor is concentrated in a few well-defined domains within the nucleoplasm of pinealocytes. Dispersion of the nuclear CREB1 increases in the main PG cell type during the dark phase, in the presence of neural norepinephrine (NE). Chronic sympathetic disruption caused by bilateral SCGx generates a relatively higher dispersion of nuclear CREB1 in pinealocytes, as compared to the SHAM group. (B) Highly dense distribution of nuclear CREB1 in non-pinealocyte cells, which was not obviously affected by temporal cues, nor by the surgical procedures. Nu: nucleus; SCGx: superior cervical ganglionectomy; ZT: *Zeitgeber* time.

Further analysis is required to define the precise moment of acquisition of the pinealocyte-specific CREB1 pattern during PG ontogeny (35), and its correlation or not with the establishment of the pineal-defining transcriptome, which occurs prior to 5 days after birth in rat (29), and to characterize the spatio-temporal patterns of nuclear CREB1 and pSer^133^-CREB1 for both the alpha- and the beta-pinealocyte identities. These two pinealocyte subpopulations differ in their transcriptomes, and in the efficiency of the molecular machinery responsible for converting the precursor N-acetylserotonin (NAS) into melatonin (30). Taken together, the pineal gland itself might be a good option for use as a circadian model to study the regulatory complexity behind transcriptional architecture and the nuclear landscape.

## Acknowledgments

We thank Julieta Scelta for technical assistance, and Raymond Astrue for editing the manuscript.

## Author contributions

LEFA and EMM designed the experiments. LEFA implemented the surgical procedures. LEFA, EV and CLF performed the experiments and collected data. LEFA and JEI developed the multi-step image processing method used to study CREB1 spatio-temporal dynamics within individual pinealocyte nuclei. LEFA, EV, CLF, JEI, MEG and EMM analyzed data. EMM provided the resources. LEFA, EV, CLF and EMM contributed to the original draft. EMM wrote the final version of the manuscript. MEG and EMM reviewed and edited the final version of the manuscript. All authors have read and agreed to the published version of the manuscript.

