## Supplemental figures for "Spatio-temporal dynamics of nuclear CREB1: what does it mean?"

### Supporting information

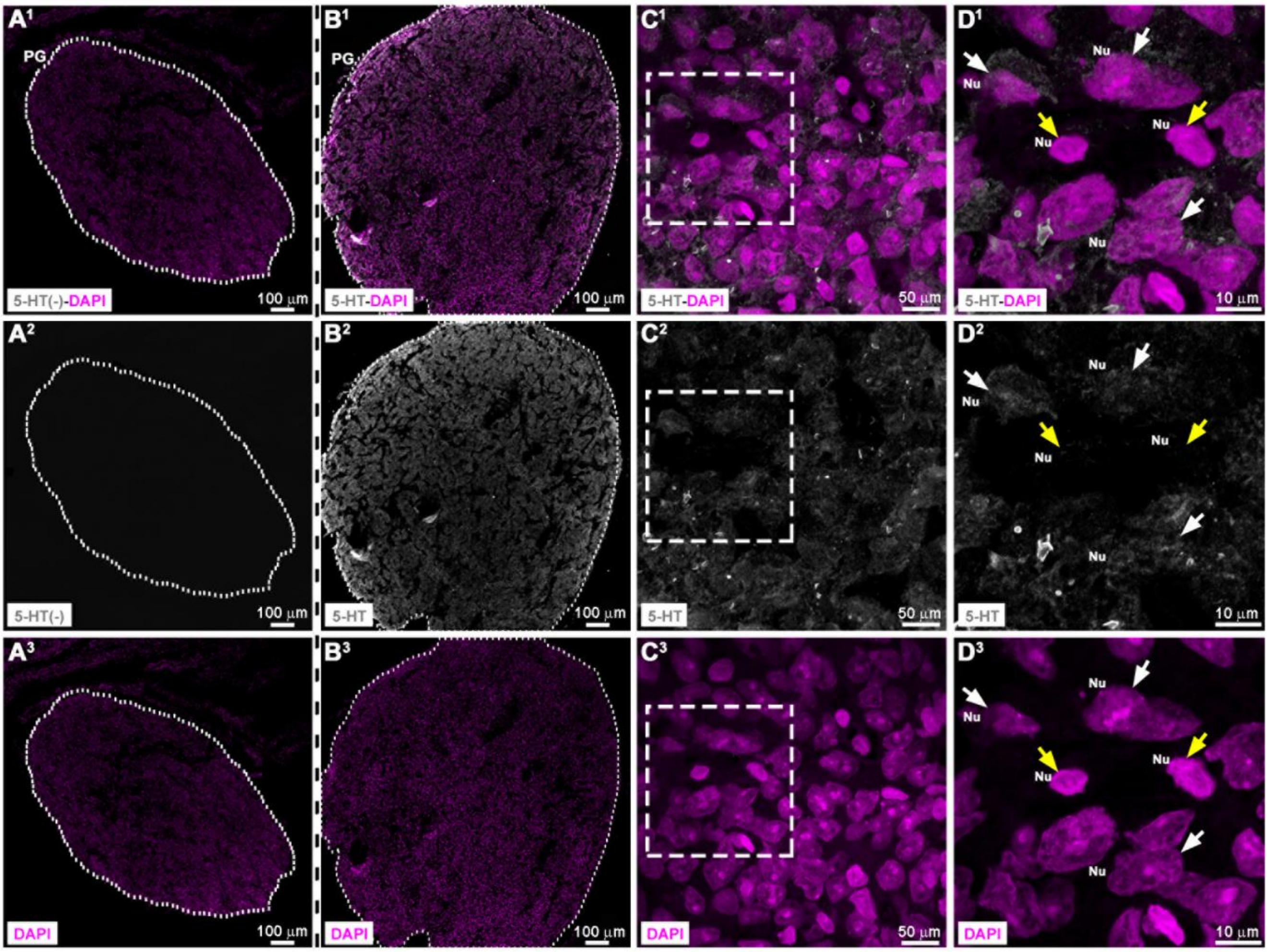


**S1 Fig. Identification of pinealocytes within the rat pineal gland.** Section of adult rat pineal gland (PG) collected at ZT6 and immunolabeled for the melatonin precursor, serotonin or 5-hydroxytryptamine (5-HT; grey). Nuclei (Nu) were dyed with 4′,6-diamidino-2-phenylindole (DAPI; magenta). ($A^1$-$A^3$) Negative control by omission of the anti-5-HT antibody. ($B^1$-$D^3$) PG section in the presence of the primary antibody and/or the nuclear dye.  ($A^1$-$B^3$) 10x images; scale bar: 100 µm. The PG perimeter is defined by a dashed white line. ($C^1$-$C^3$) 2x digital zooms from 60x images; scale bar: 50 µm. ($D^1$-$D^3$) 2.5x digital zooms from 100x images of the insets shown in $C^1$-$C^3$; scale bar: 10 µm. Predominant pinealocytes are easily identified due to their nuclear architecture and their immunoreactivity for 5-HT (white arrows). Non-pinealocyte cells with homogenous and compact chromatin showed no signal for 5-HT (yellow arrows).

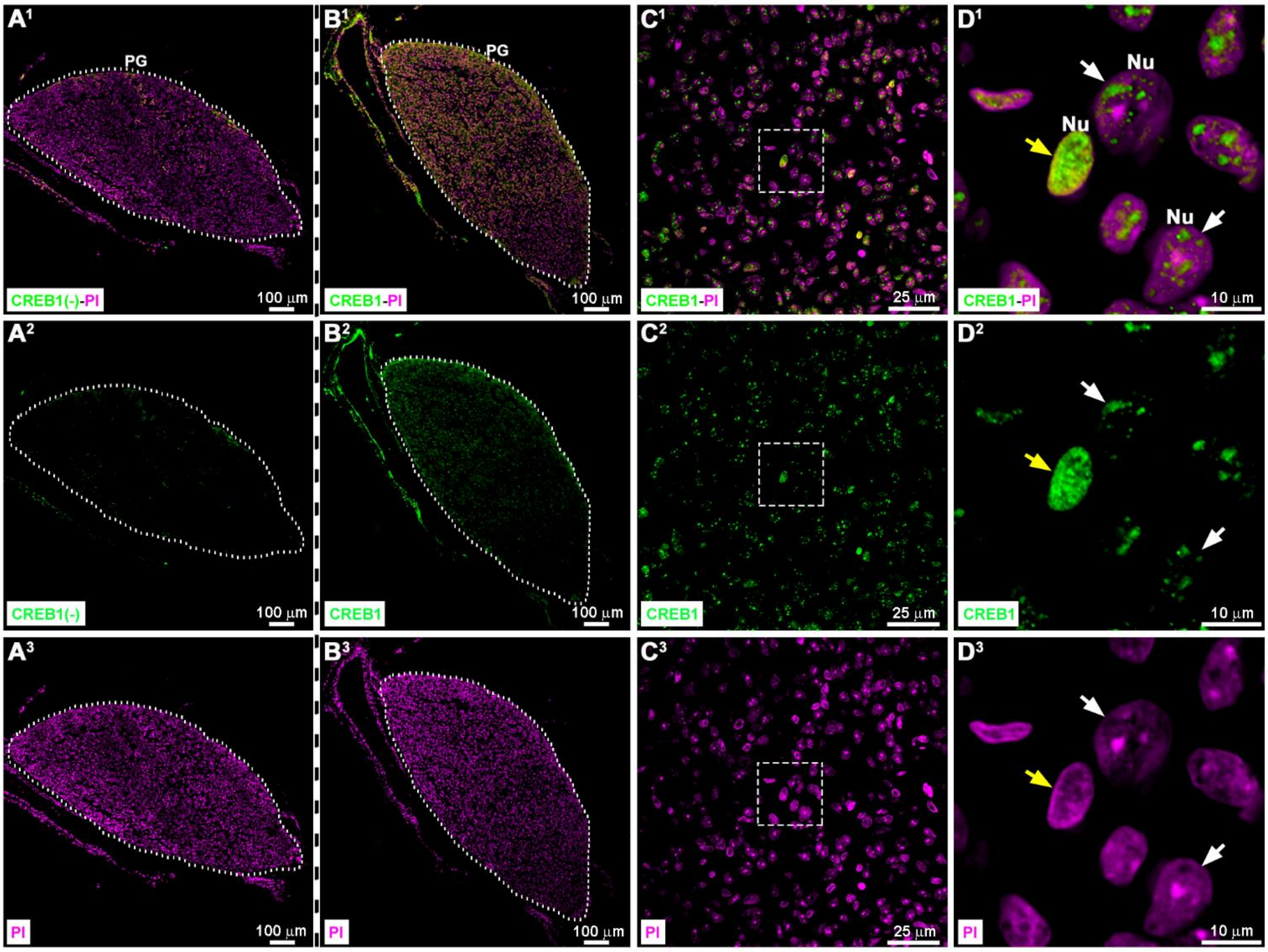


**S2 Fig. Heterogeneous distribution of CREB1 among the different cell types within the rat pineal gland.** Section of an adult pineal gland (PG) collected at ZT14 and immunolabeled for CREB1 (green). Cell nuclei (Nu) were stained with propidium iodide (PI; magenta). ($A^1$-$A^3$) Negative control by omission of the anti-CREB1 antibody. ($B^1$-$D^3$) PG section in the presence of the primary antibody and/or the nuclear dye. ($A^1$-$B^3$) 10x images; scale bar: 100 µm. The PG perimeter is defined by a dashed white line. ($C^1$-$C^3$) 60x images; scale bar: 25 µm. ($D^1$-$D^3$) 5x digital zooms of the insets shown in $C^1$-$C^3$; scale bar: 10 µm. CREB1 occupies discrete domains within the nuclei of pinealocytes (white arrows). In contrast, CREB1 is compactly and homogenously distributed in the nucleus of a non-pinealocyte cell (yellow arrow). ZT: *Zeitgeber* time.

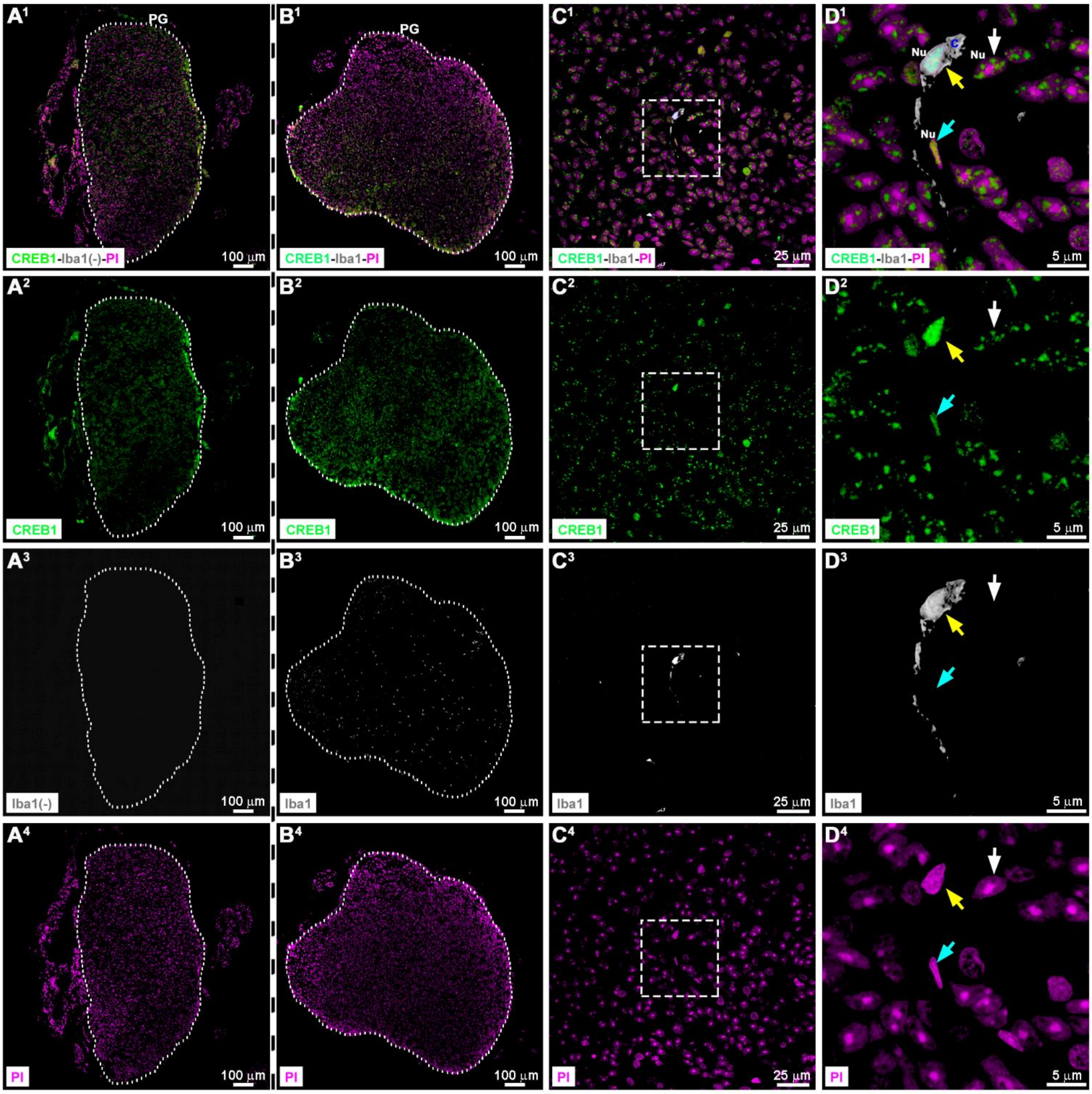


**S3 Fig. Homogeneous and dense distribution of CREB1 in the nuclei of phagocytes within the rat pineal gland.** Sections of adult pineal glands (PG) immunolabeled for CREB1 (green) and the microglia/macrophage-specific ionized calcium-binding adapter molecule 1 (Iba1; grey). Nuclei (Nu) were dyed with propidium iodide (PI; magenta). ($A^1$-$A^4$) Negative control in the absence of the anti-Iba1 antibody. ($B^1$-$D^4$) Double immunolabeling of a PG section in the presence of the nuclear dye. Single channels and merged images are shown. ($A^1$-$B^4$) 10x images; scale bar: 100 µm. The PG perimeter is defined by a dashed white line. ($C^1$-$C^4$) 60x images; scale bar: 25 µm. ($D^1$-$D^4$) 2x digital zooms of the insets shown in $C^1$-$C^4$; scale bar: 5 µm. CREB1 is present in the nucleus of an Iba1$^+$ cell, with a homogeneous and dense distribution (yellow arrow). A similar compact pattern of CREB1 is observed in an elongated nucleus of a non-pinealocyte cell (cyan arrow). A pinealocyte nucleus with discrete domains of CREB1 is indicated (white arrow). C: cytoplasm.

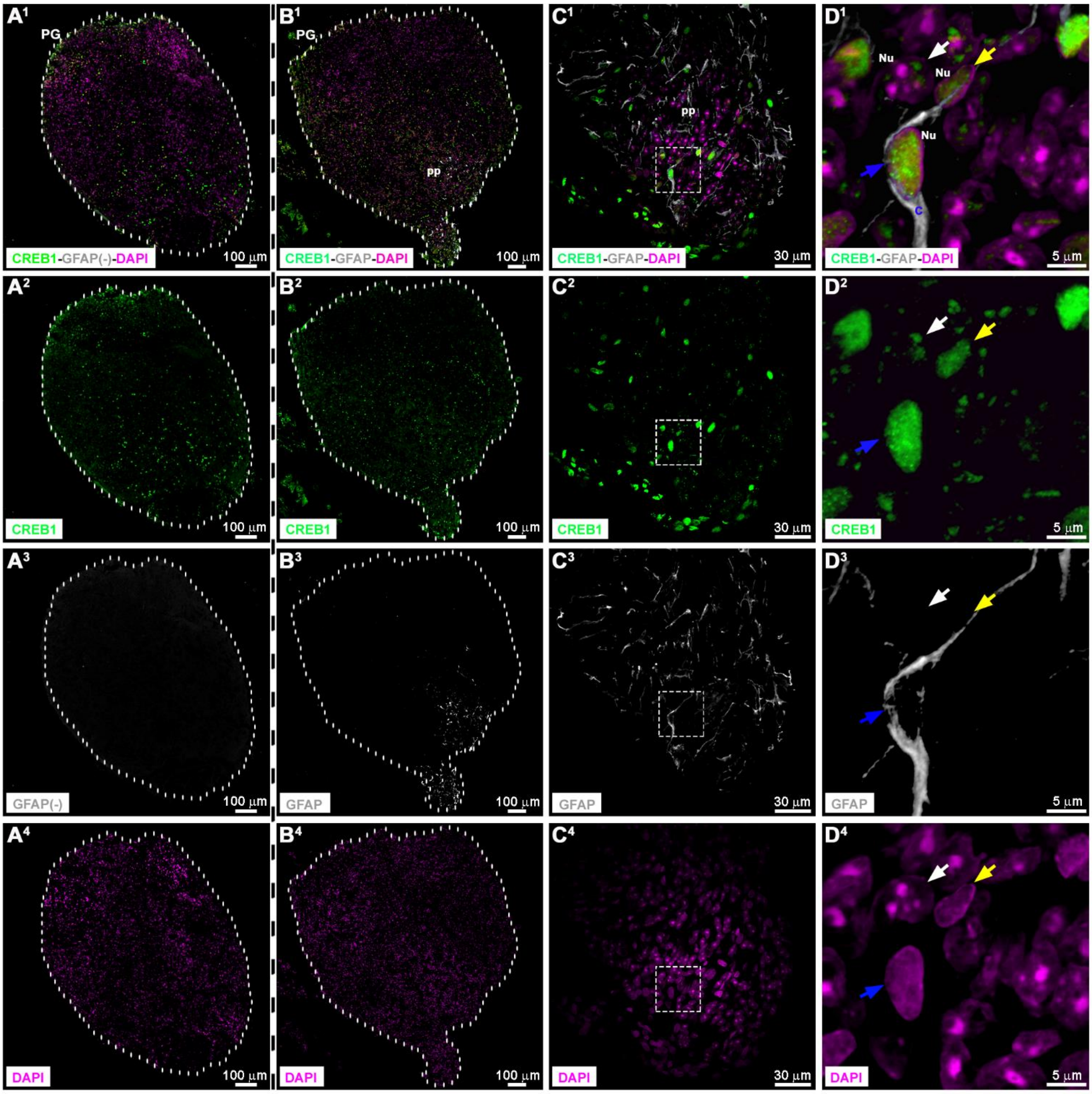


**S4 Fig. Homogeneous and dense distribution of CREB1 in the nuclei of astrocytes within the rat pineal gland.** Sections of adult pineal glands (PG) immunolabeled for CREB1 (green), and the glial fibrillary acidic protein (GFAP; grey) which is a marker commonly used for the identification of astrocytes and precursor-like cells. Nuclei (Nu) were dyed with 4′,6-diamidino-2-phenylindole (DAPI; magenta). ($A^1$-$A^4$) Negative control by the omission of the anti-GFAP antibody. ($B^1$-$D^4$) Double immunolabeling of a PG section in the presence of DAPI. Single channels and merged images are shown. ($A^1$-$B^4$) 10x images; scale bar: 100 µm. The PG perimeter is defined by a dashed white line. ($C^1$-$C^4$) 60x images; scale bar: 30 µm. ($D^1$-$D^4$) 6.1x digital zooms of the insets shown in $C^1$-$C^4$; scale bar: 5 µm. In the proximal pole (pp) of the PG section, CREB1 is present in the nuclei of $GFAP^+$ cells, with a homogeneous and dense distribution (blue arrow). A similar compact pattern of CREB1 is observed in a non-pinealocyte cell, negative for GFAP (yellow arrow). A pinealocyte nucleus with discrete domains of CREB1 is indicated (white arrow). C: cytoplasm.

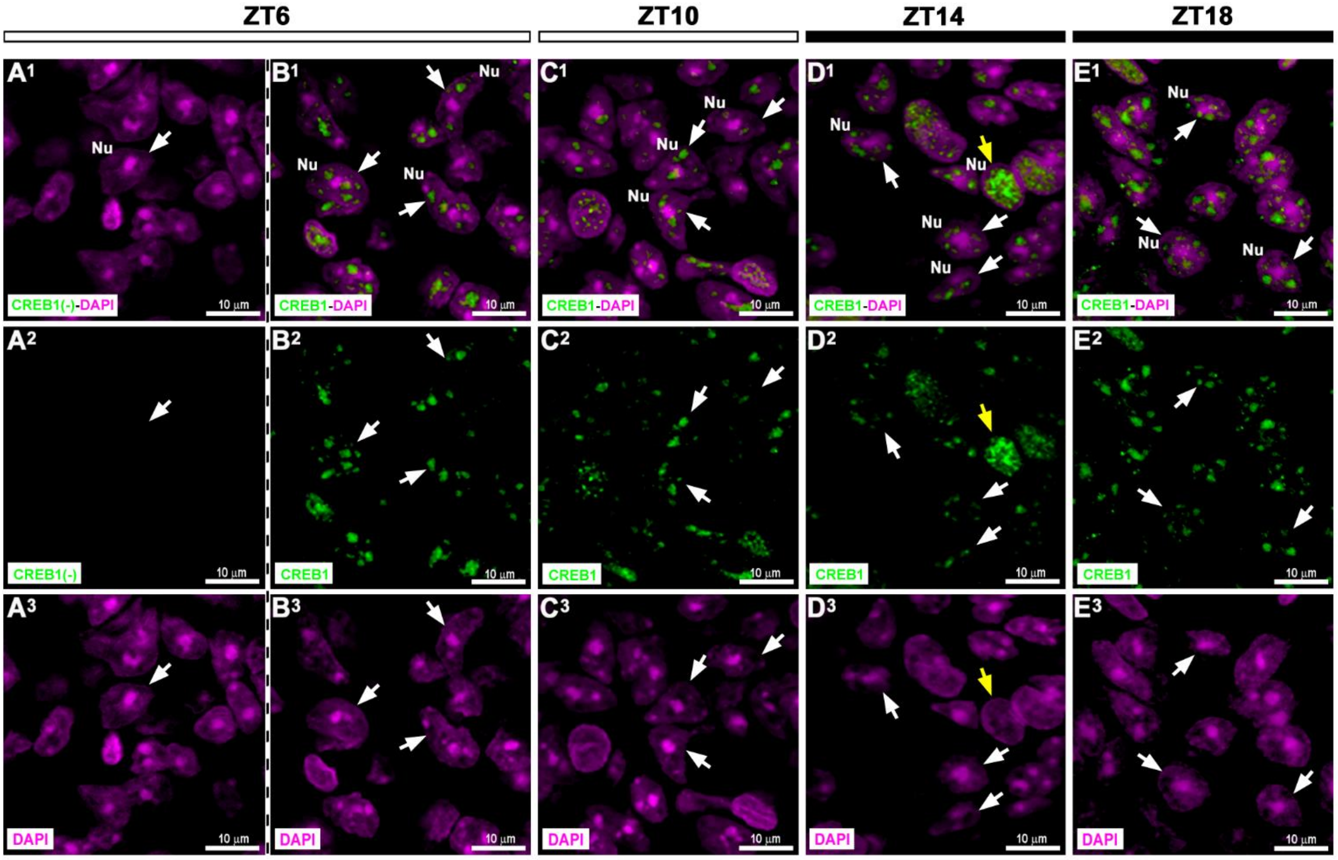


**S5 Fig. Heterogenous distribution of CREB1 among rat pinealocyte nuclei.** Sections of adult rat pineal glands (PG) immunolabeled for CREB1 (green). Nuclei (Nu) were stained with 4′,6-diamidino-2-phenylindole (DAPI; magenta). ($A^1$-$A^3$) Incubation without the anti-CREB1 antibody (negative control). ($B^1$-$E^3$) Immunostaining for CREB1 at daytime (ZT6 and ZT10), and at nighttime (ZT14 and ZT18). ($A^1$-$E^3$) 2x digital zooms from 60x images; scale bar: 10 µm. Heterogeneity in CREB1 distribution is observed among pinealocyte nuclei (white arrows) at a defined ZT and among ZTs. A non-pinealocyte nucleus, densely immunoreactive for CREB1, is indicated (yellow arrow). ZT: *Zeitgeber* time.

# ZT6

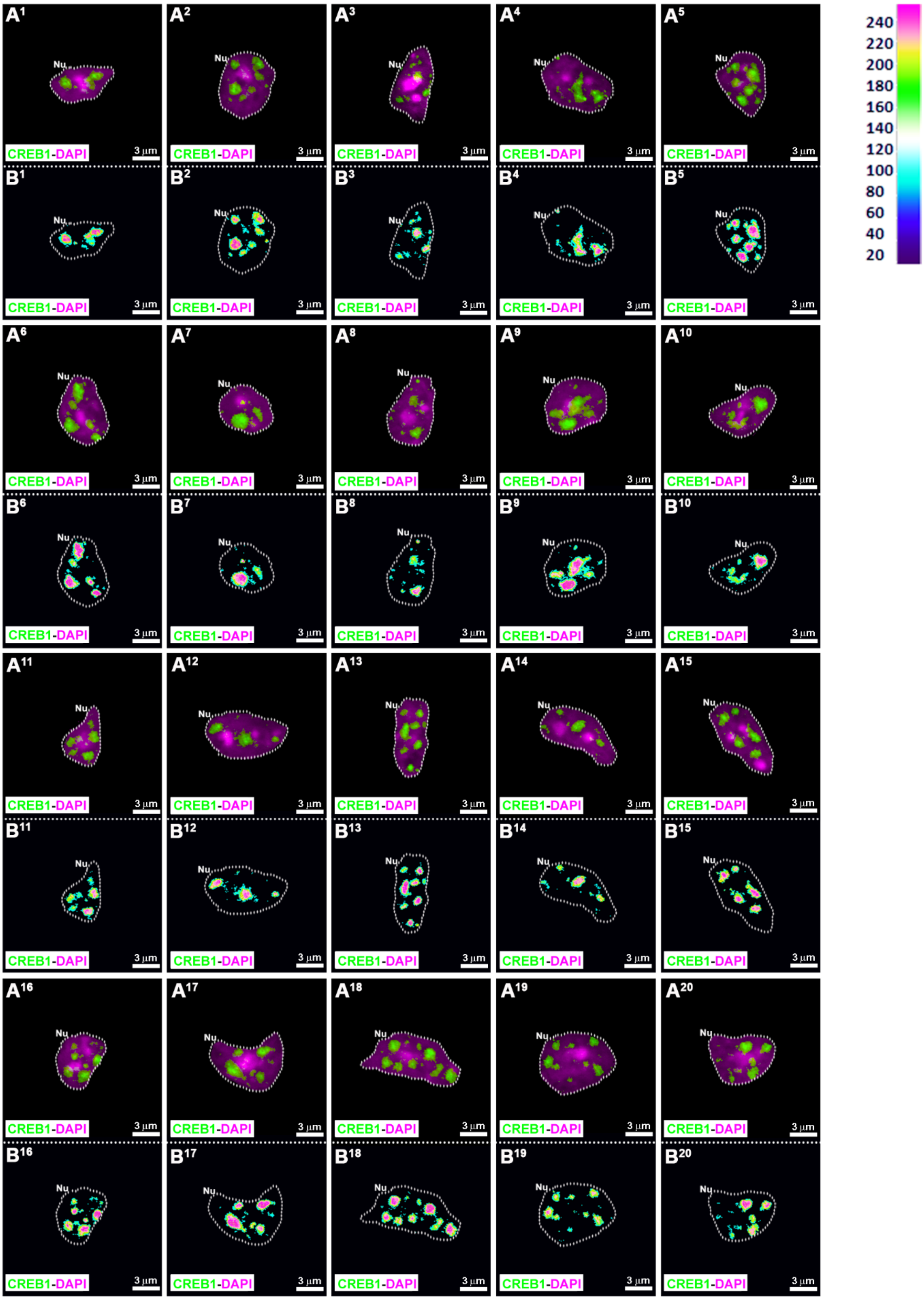
A1 A2 A3 A4 A5
B1 B2 B3 B4 B5
A6 A7 A8 A9 A10
B6 B7 B8 B9 B10
A11 A12 A13 A14 A15
B11 B12 B13 B14 B15
A16 A17 A18 A19 A20
B16 B17 B18 B19 B20
Nu
CREB1-DAPI
3 µm
240
220
200
180
160
140
120
100
80
60
40
20

**S6 Fig . Spatial distribution of CREB1 within individual pinealocyte nuclei at ZT6.** ($A^{1}$-$A^{20}$) Twenty representative pinealocyte nuclei (Nu), immunolabeled for CREB1 (green), were isolated from pineal gland sections of adult rats sacrificed at ZT6. Nuclei were counterstained with 4′,6-diamidino-2-phenylindole (DAPI; magenta). ($B^{1}$-$B^{20}$) Schematic representations of the fluorescence intensity of CREB1 for each pixel within the nuclei shown in $A^{1}$-$A^{20}$. The fluorescence intensity ranges from 0 to 255. To build these representations, the intensity values were look-up table (LUT) mapped to color values using the ImageJ software (Version 1.52d, NIH, USA). ($A^{1}$-$B^{20}$) 2x digital zooms from 60x images; scale bar: 3 µm. Nuclei were selected from 4 pineal glands (PG). The nuclear perimeter is defined by a dashed white line. ZT: *Zeitgeber* time.

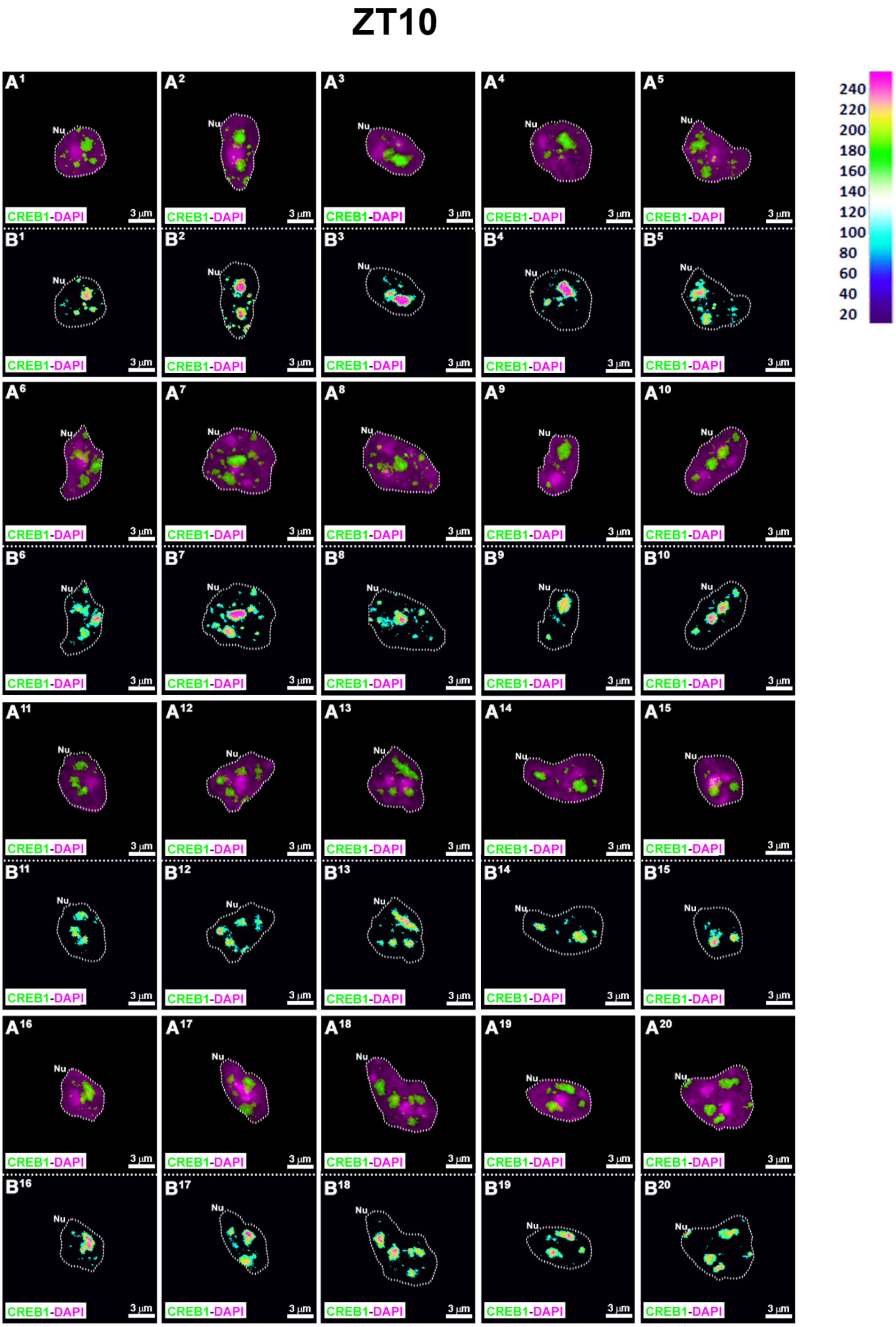
ZT10
A1
A2
A3
A4
A5
B1
B2
B3
B4
B5
A6
A7
A8
A9
A10
B6
B7
B8
B9
B10
A11
A12
A13
A14
A15
B11
B12
B13
B14
B15
A16
A17
A18
A19
A20
B16
B17
B18
B19
B20
Nu
CREB1-DAPI
3 µm
240
220
200
180
160
140
120
100
80
60
40
20

**S7 Fig. Spatial distribution of CREB1 within individual pinealocyte nuclei at ZT10.** ($A^1$-$A^{20}$) Twenty representative pinealocyte nuclei (Nu), immunolabeled for CREB1 (green), were isolated from pineal gland sections of adult rats sacrificed at ZT10. Nuclei were counterstained with 4′,6-diamidino-2-phenylindole (DAPI; magenta). ($B^1$-$B^{20}$) Schematic representations of the fluorescence intensity of CREB1 for each pixel within the nuclei shown in $A^1$-$A^{20}$. The fluorescence intensity ranges from 0 to 255. ($A^1$-$B^{20}$) 2x digital zooms from 60x images; scale bar: 3 µm. Nuclei were selected from 4 pineal glands (PG). The nuclear perimeter is defined by a dashed white line. ZT: *Zeitgeber* time.

ZT14

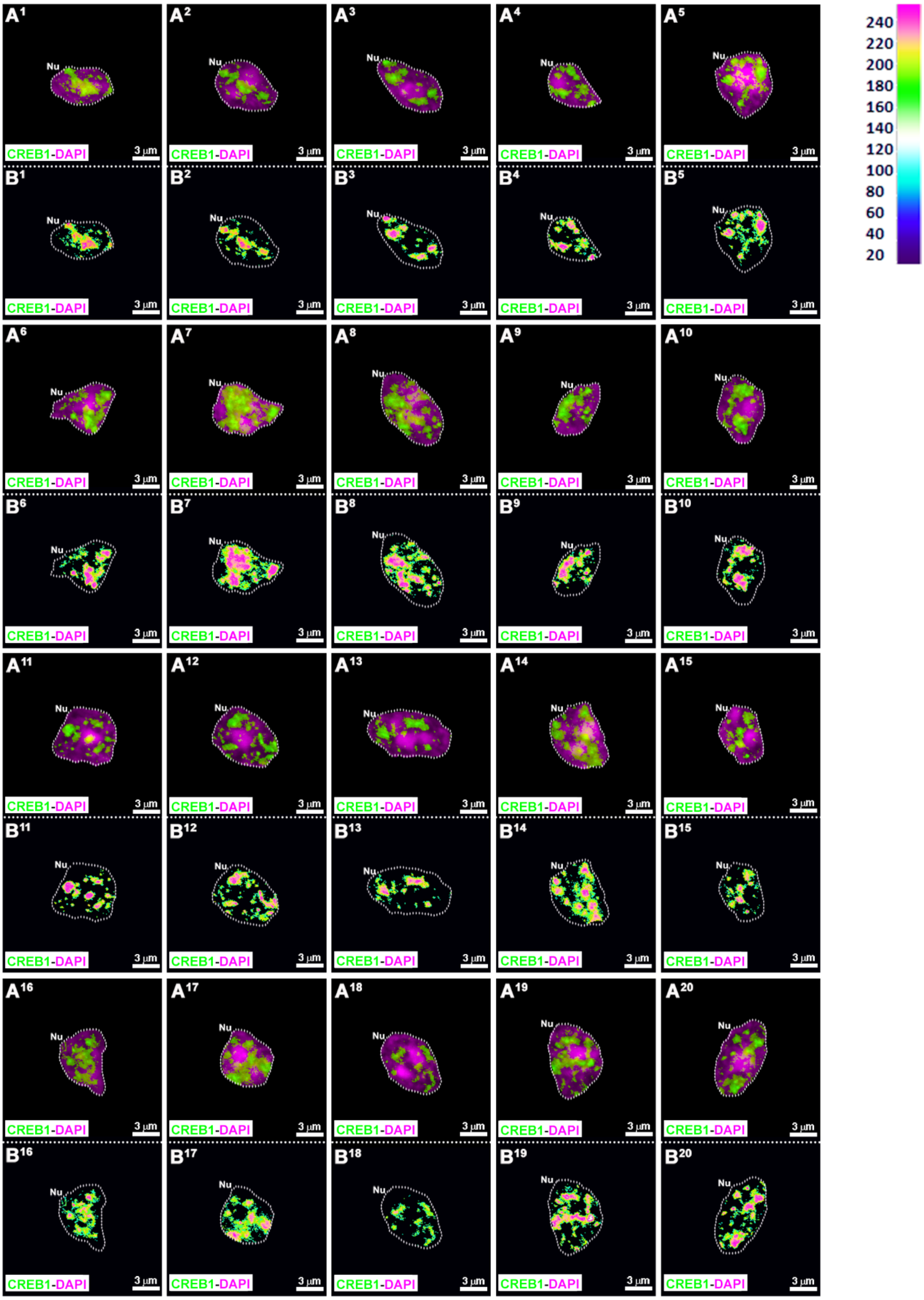

A1
A2
A3
A4
A5
B1
B2
B3
B4
B5
A6
A7
A8
A9
A10
B6
B7
B8
B9
B10
A11
A12
A13
A14
A15
B11
B12
B13
B14
B15
A16
A17
A18
A19
A20
B16
B17
B18
B19
B20
Nu
CREB1-DAPI
3 µm
240
220
200
180
160
140
120
100
80
60
40
20

**S8 Fig. Spatial distribution of CREB1 within individual pinealocyte nuclei at ZT14.** ($A^1$-$A^{20}$) Twenty representative pinealocyte nuclei (Nu), immunolabeled for CREB1 (green), were isolated from pineal gland sections of adult rats sacrificed at ZT14. Nuclei were counterstained with 4′,6-diamidino-2-phenylindole (DAPI; magenta). ($B^1$-$B^{20}$) Schematic representations of the fluorescence intensity of CREB1 for each pixel within the nuclei shown in $A^1$-$A^{20}$. The fluorescence intensity ranges from 0 to 255. ($A^1$-$B^{20}$) 2x digital zooms from 60x images; scale bar: 3 μm. Nuclei were selected from 3 pineal glands (PG). The nuclear perimeter is defined by a dashed white line. ZT: *Zeitgeber* time.

# ZT18

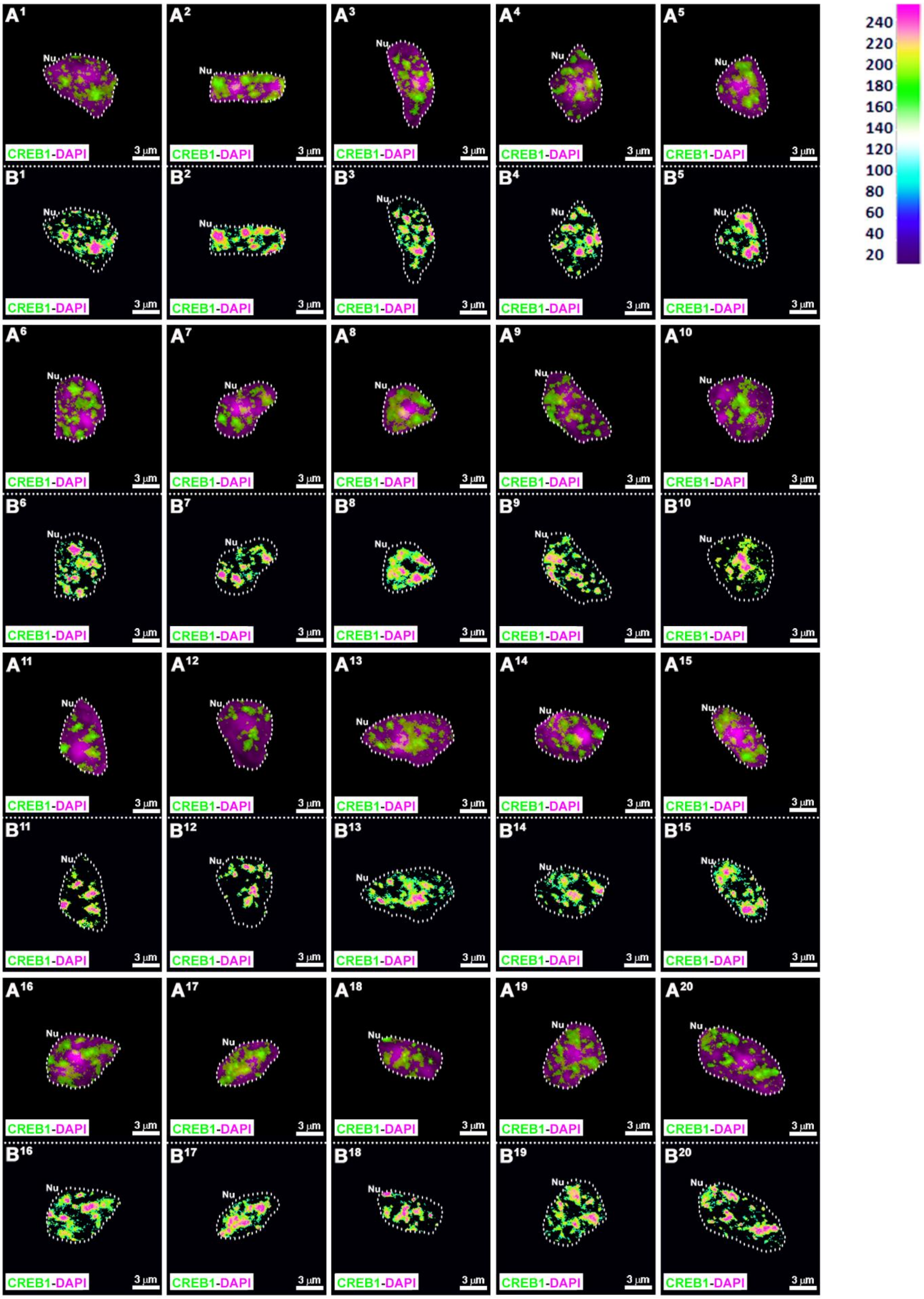
A1 A2 A3 A4 A5 A6 A7 A8 A9 A10 A11 A12 A13 A14 A15 A16 A17 A18 A19 A20
B1 B2 B3 B4 B5 B6 B7 B8 B9 B10 B11 B12 B13 B14 B15 B16 B17 B18 B19 B20
Nu
CREB1-DAPI
3 µm
240
220
200
180
160
140
120
100
80
60
40
20

**S9 Fig. Spatial distribution of CREB1 within individual pinealocyte nuclei at ZT18.** ($A^1$-$A^{20}$) Twenty representative pinealocyte nuclei (Nu), immunolabeled for CREB1 (green), were isolated from pineal gland sections of adult rats sacrificed at ZT18. Nuclei were counterstained with 4′,6-diamidino-2-phenylindole (DAPI; magenta). ($B^1$-$B^{20}$) Schematic representations of the fluorescence intensity of CREB1 for each pixel within the nuclei shown in $A^1$-$A^{20}$. The fluorescence intensity ranges from 0 to 255. ($A^1$-$B^{20}$) 2x digital zooms from 60x images; scale bar: 3 μm. Nuclei were selected from 3 pineal glands (PG). The nuclear perimeter is defined by a dashed white line. ZT: *Zeitgeber* time.

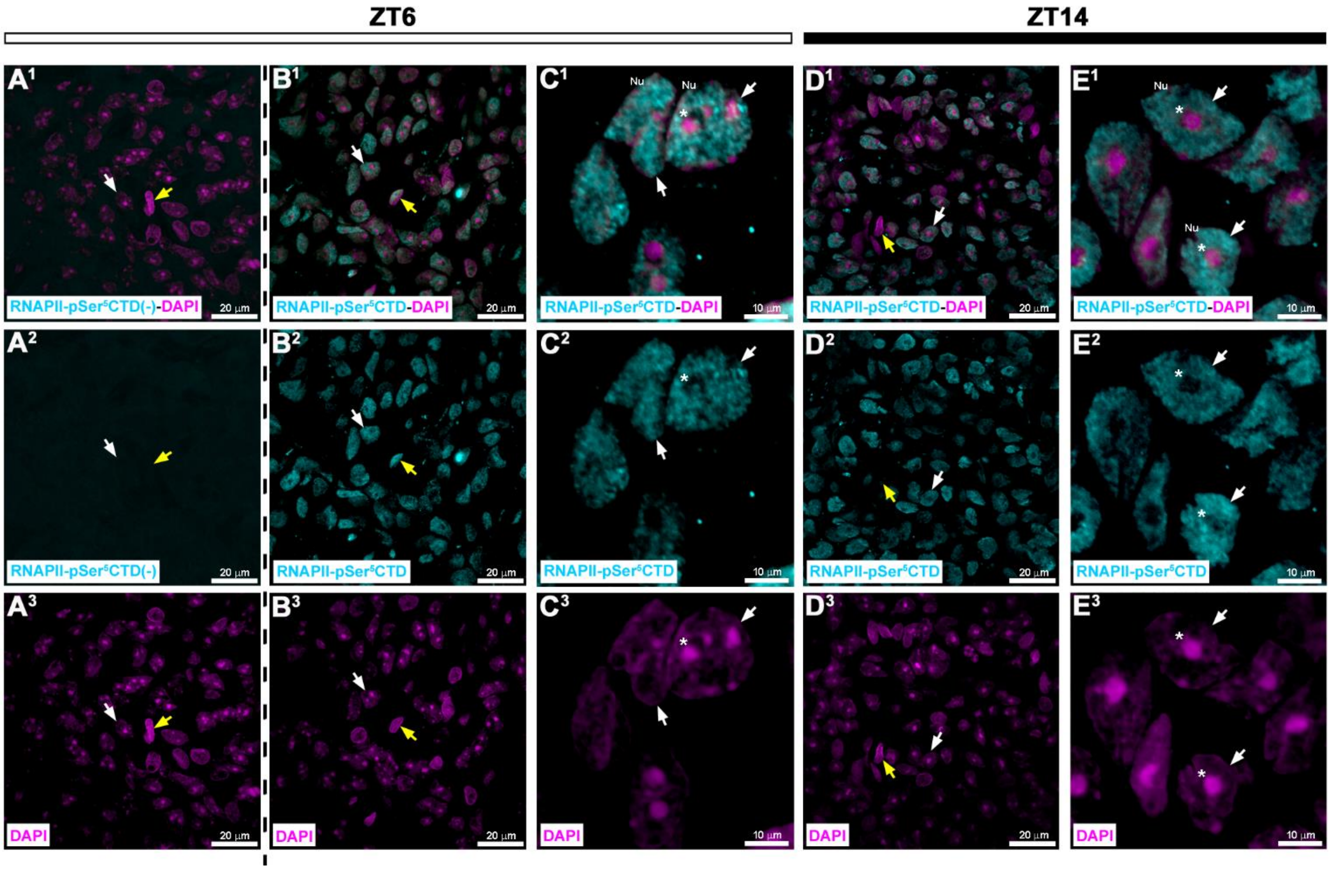


**S10 Fig. Day/night analysis of the nuclear distribution of the phosphorylated RNA polymerase II in the rat pineal gland.** Sections of adult rat pineal glands collected during the daytime (ZT6) and nighttime (ZT14), immunolabeled for a particular phosphorylated form of the RNA polymerase II (RNAPII-pSer$^5$CTD; cyan). Nuclei (Nu) were counterstained with 4′,6-diamidino-2-phenylindole (DAPI; magenta). (A$^1$-A$^3$) Negative control by omission of the anti-RNAPII-pSer$^5$CTD antibody. (B$^1$-E$^3$) Single channels and merged images are shown. (A$^1$-A$^3$, B$^1$-B$^3$, D$^1$-D$^3$) 60x images; scale bar: 20 µm. (C$^1$-C$^3$, E$^1$-E$^3$) 10x digital zooms from 60x images; scale bar: 5 µm. Pinealocyte nuclei immunoreactive for RNAPII-pSer$^5$CTD are indicated (white arrows). Non-pinealocyte nuclei with different levels of RNAPII-pSer$^5$CTD are also indicated (yellow arrows). Asterisks: nucleoli without RNAPII-pSer$^5$CTD signal, present within pinealocyte nuclei. CTD: C-terminal repeat domain (YSPTSPS) of the RNA polymerase II.

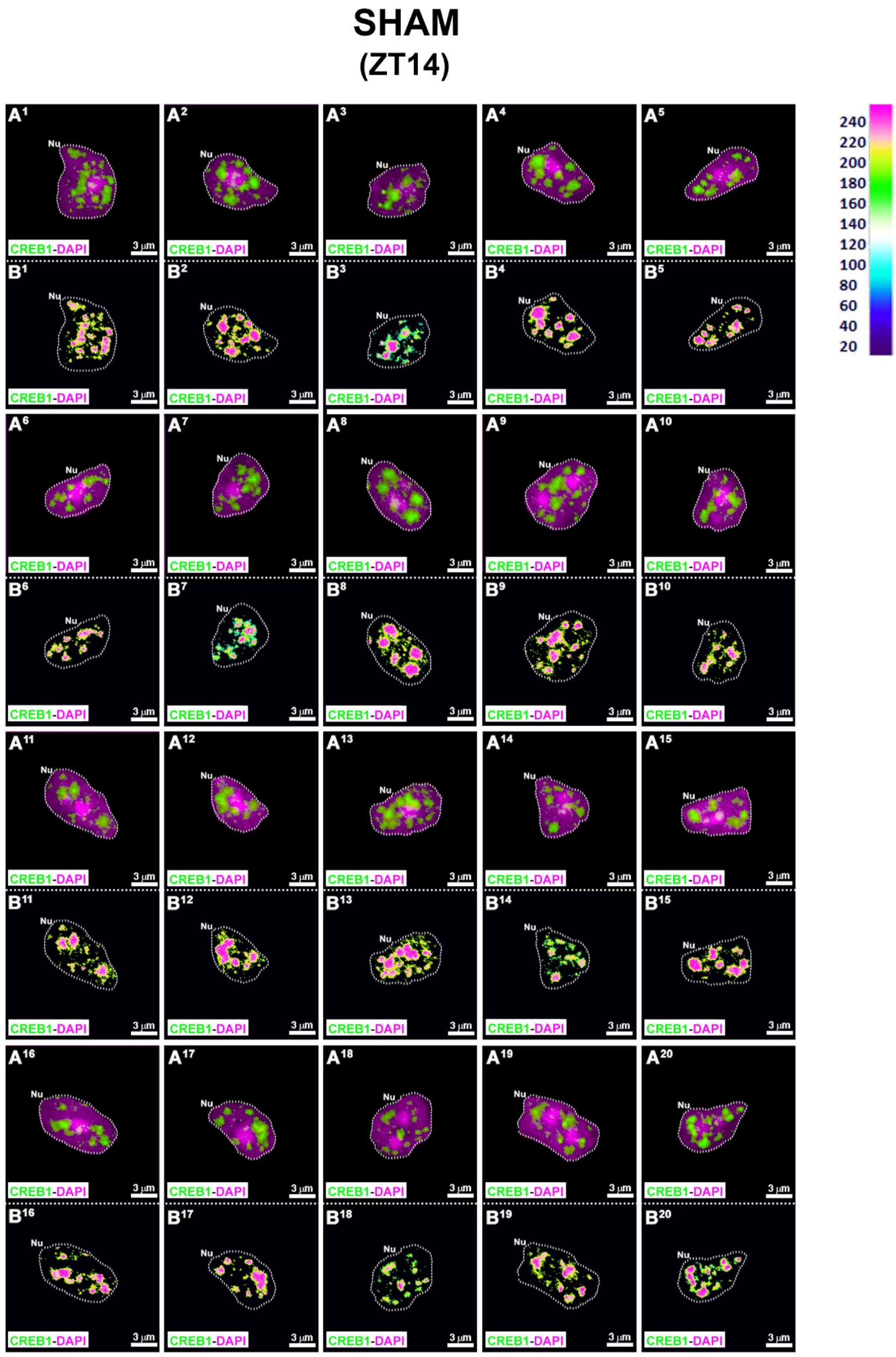
SHAM
(ZT14)
A1 A2 A3 A4 A5 A6 A7 A8 A9 A10 A11 A12 A13 A14 A15 A16 A17 A18 A19 A20
B1 B2 B3 B4 B5 B6 B7 B8 B9 B10 B11 B12 B13 B14 B15 B16 B17 B18 B19 B20
Nu
CREB1-DAPI
3 μm
240
220
200
180
160
140
120
100
80
60
40
20

**S11 Fig. Spatial distribution of CREB1 within individual pinealocyte nuclei of adult sham-operated rats.** ($A^{1}$-$A^{20}$) Twenty representative pinealocyte nuclei (Nu), immunoreactive for CREB1 (green), were isolated from pineal gland sections of adult sham-operated rats sacrificed three weeks after surgery at ZT14 (N=4). Nuclei were dyed with 4′,6-diamidino-2-phenylindole (DAPI; magenta). The fluorescence intensity ranges from 0 to 255. ($B^{1}$-$B^{20}$) Schematic representations of the fluorescence intensity of CREB1 for each pixel within the nuclei shown in $A^{1}$-$A^{20}$. ($A^{1}$-$B^{20}$) 2x digital zooms from 60x images; scale bar: 3 μm. The nuclear perimeter is defined by a dashed white line. ZT: *Zeitgeber* time.

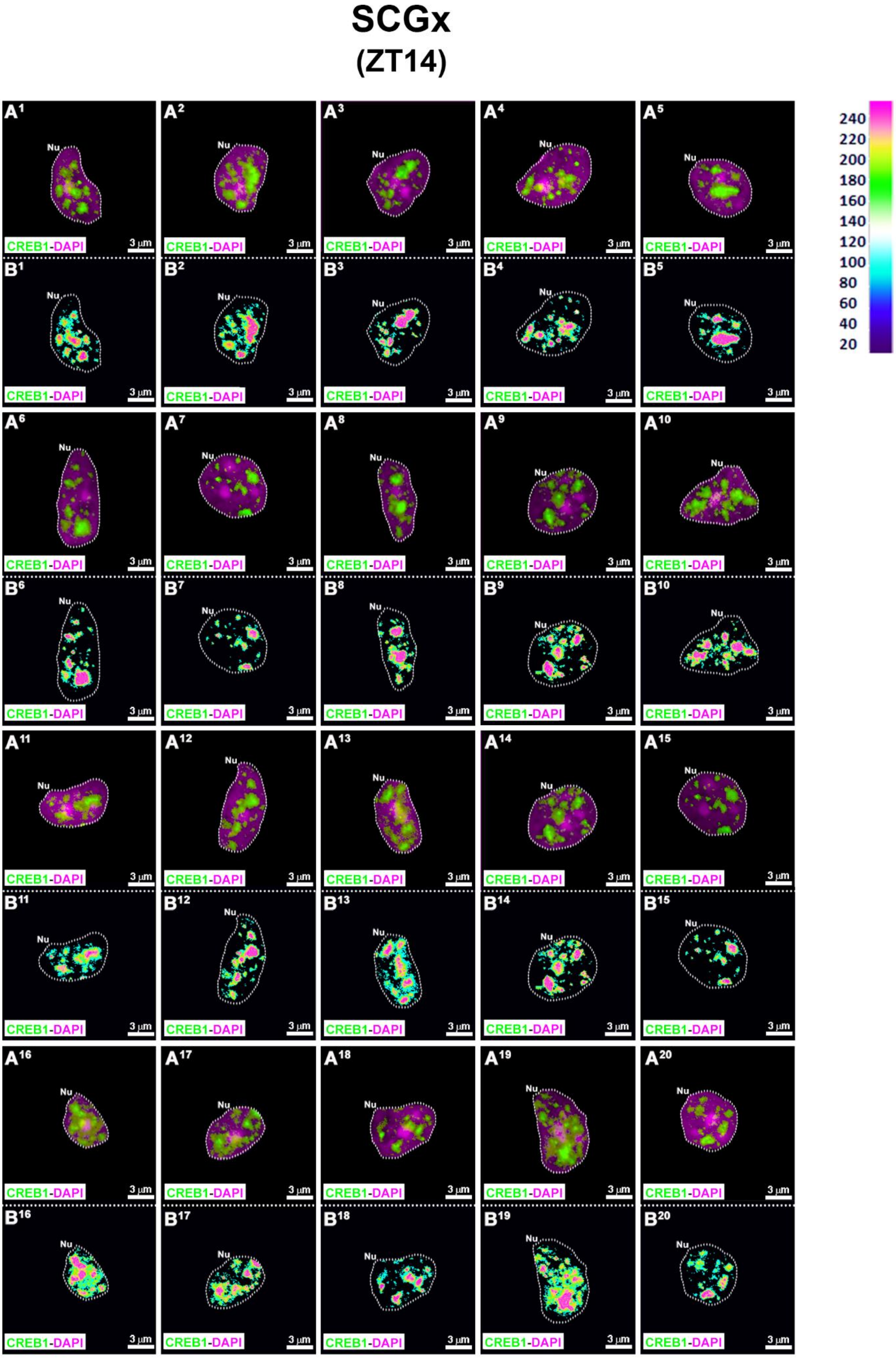
SCGx
(ZT14)
A1 A2 A3 A4 A5 A6 A7 A8 A9 A10 A11 A12 A13 A14 A15 A16 A17 A18 A19 A20
B1 B2 B3 B4 B5 B6 B7 B8 B9 B10 B11 B12 B13 B14 B15 B16 B17 B18 B19 B20
Nu
CREB1-DAPI
3 μm
240
220
200
180
160
140
120
100
80
60
40
20

**S12 Fig. Spatial distribution of CREB1 within individual pinealocyte nuclei at ZT14, after chronic bilateral superior cervical ganglionectomy.** Spatial distribution of CREB1 within individual pinealocyte nuclei at ZT14, after chronic bilateral superior cervical ganglionectomy. ($A^1$-$A^{20}$) Twenty representative pinealocyte nuclei (Nu), immunoreactive for CREB1 (green), were isolated from pineal gland sections of adult SCGx rats sacrificed three weeks after surgery at ZT14 (N=4). Nuclei were dyed with 4′,6-diamidino-2-phenylindole (DAPI; magenta). (B1-$B^{20}$) Schematic representations of the fluorescence intensity of CREB1 for each pixel within the nuclei shown in $A^1$-$A^{20}$. The fluorescence intensity ranges from 0 to 255. ($A^1$-$B^{20}$) 2x digital zooms from 60x images; scale bar: 3 μm. The nuclear perimeter is defined by a dashed white line. SCGx: superior cervical ganglionectomy; ZT: Zeitgeber time.
